# FGF signaling in development beyond canonical pathways

**DOI:** 10.1101/2020.05.13.093252

**Authors:** Ayan T. Ray, Pierre Mazot, J. Richard Brewer, Catarina Catela, Colin J. Dinsmore, Philippe Soriano

**Affiliations:** Department of Cell, Developmental, and Regenerative Biology Icahn School of Medicine at Mount Sinai, New York, NY 10029

**Keywords:** FGF, ERK1/2, Craniofacial development, Neural crest, Cell adhesion

## Abstract

FGFs are key developmental regulators which engage a signal transduction cascade through receptor tyrosine kinases, typically involving ERK1/2, PI3K/AKT, and other effectors. However, it remains unknown if all FGF activities depend on kinase activity or these canonical signal transduction cascades. To address these questions, we generated allelic series of knock-in *Fgfr1* and *Fgfr2* mouse strains, carrying point mutations that disrupt binding of signaling effectors to the receptors, alone or in combination. We also produced a kinase dead allele of *Fgfr2* which broadly phenocopies the null mutant. When interrogated in cranial neural crest cells, point mutations in either receptor revealed discrete functions for signaling pathways in specific craniofacial contexts, but failed to recapitulate the single or double null mutant phenotypes even in their most extensive combination. Furthermore, we found that together these signaling mutations abrogated the established FGF-induced signal transduction pathways, yet certain FGF functions such as cell-matrix and cell-cell adhesion remained unaffected. Our studies establish combinatorial roles of both *Fgfr1* and *Fgfr2* in development and identify novel kinase-dependent cell adhesion properties of FGF receptors, independent of well-established roles in intracellular signaling.

---

Classic models of receptor tyrosine kinase activation involve ligand binding, receptor dimerization, transactivation of the kinase domain, phosphorylation of intracellular tyrosines, and binding of effectors that orchestrate activation of downstream signaling pathways ^1^. Different thresholds in dimer strength and stability may also come into play to regulate signaling dynamics ^2–4^. For FGFRs, where downstream pathways have been particularly well studied, numerous lines of evidence point to ERK1/2 as the main effector of FGF signaling ^5,6^. Although characterization of effector binding to RTKs provides critical insights on signaling specificity, assessing relative pathway significance requires *in vivo* validation. A previous analysis showed that knock-in *Fgfr1* point mutations disrupting binding of multiple signaling effectors, alone or in combination, did not recapitulate the *Fgfr1*^−/−^ phenotype despite eliminating ERK1/2 outputs ^7^. These results, although suggestive of additional signaling pathways, were difficult to interpret due to expression of another receptor, FGFR2. Using cranial neural crest (cNCC)-derived mesenchyme to understand FGF signaling, we now investigate the relative contributions of both FGF receptors and their signaling outputs to craniofacial development. We find that signaling mutations for each receptor disrupt classical signal transduction pathways in the absence of the other receptor, but do not recapitulate the null phenotypes. We furthermore show that FGF signaling is kinase-dependent and identify non-canonical functions of FGF signaling that help reconcile the gap in our phenotypic analyses.

## *Fgfr1* and *Fgfr2* orchestrate craniofacial development by regulating cell survival

To interrogate FGFR1/2 functions, we initially investigated how loss of both receptors in cNCCs influences craniofacial development. We found both receptors extensively co-expressed in the cNCC-derived mesenchyme and overlying epithelia, using fluorescent *Fgfr1* and *Fgfr2* reporter alleles (Extended Data Fig. 1A). Next, conditional null alleles (henceforth denoted *cKO*) of *Fgfr1* and *Fgfr2* were combined with *Wnt1Cre* drivers active in NCCs ^8,9^. At embryonic day (E) 18.5, *Fgfr1*^*cKO/cKO*^ embryos displayed a fully penetrant facial cleft, *Fgfr2*^*cKO/cKO*^ mutants had no overt phenotype, and *Fgfr1*^*cKO/cKO*^; *Fgfr2*^*cKO/cKO*^ double mutants exhibited severe agenesis of most NCC derived craniofacial structures including the frontal and nasal bones, nasal cartilage, maxilla, and mandible (Fig. 1A and Extended Data Fig. 1B). Both *Fgfr1*^*cKO/cKO*^; *Fgfr2*^*cKO/+*^ and *Fgfr1*^*cKO/cKO*^; *Fgfr2*^*cKO/cKO*^ embryos exhibited defects in mandible development with hypomorphic proximal structures including angular and coronoid processes (Extended Data Fig. 1C). Overall, double mutants showed reduced or no ossification of cNCC-derived craniofacial skeletal structures (Extended Data Fig. 1B-D and Supplementary Information Fig. S1A-D). At E9.5 and E10.5, *Fgfr1*^*cKO/cKO*^; *Fgfr2*^*cKO/cKO*^; *ROSA26*^*mT/mG*^ double mutants developed hypoplastic pharyngeal arches with reduced NCC lineage GFP^+^ cells (Fig. 1B and Extended Data Fig. 2A). We observed reduced expression of facial prominence markers along with midline morphogenesis defects, but ectoderm gene expression remained unaffected, suggesting reduced NCC numbers rather than patterning defects (Extended Data Fig. 2B). FGFR1 and FGFR2 are thought to function predominantly in mesenchymal or epithelial contexts, respectively, since *Fgfr1c* mutants recapitulate many aspects of the *Fgfr1*^−/−^; phenotype ^10^, and *Fgfr2b* mutants are reminiscent of *Fgfr2*^−/−^ embryos ^11^. Here, we show that both receptors function coordinately within the NCC-derived mesenchyme, as combined loss of *Fgfr1* and *Fgfr2* leads to significantly more severe midface and mandibular defects than loss of either receptor alone.

**Figure 1:**
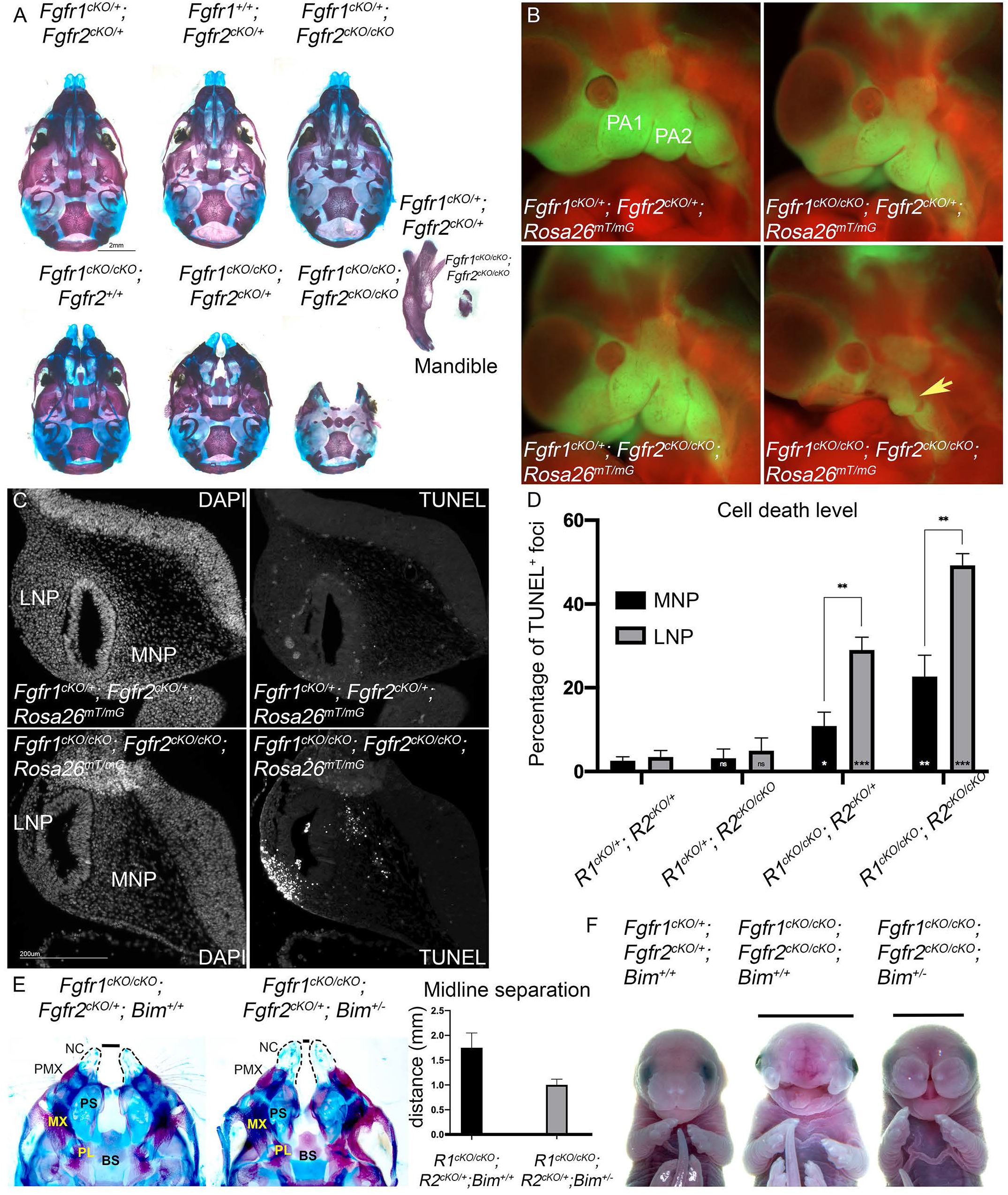
Defects in craniofacial morphogenesis in *Fgfr1/2* double mutants. (A) Alcian blue/ alizarin red staining of mouse skulls at E18.5 showed a facial cleft in *Fgfr1*^*cKO/cKO*^ embryos, which was exacerbated in *Fgfr1*^*cKO/cKO*^; *Fgfr2*^*cKO*/+^ mutants. *Fgfr1*^*cKO/cKO*^; *Fgfr2*^*cKO/cKO*^ mutants exhibited the most severe defect, with agenesis of proximal structures, as well as severe reduction of the mandible. (B) *Fgfr1*^*cKO/cKO*^; *Fgfr2*^*cKO/cKO*^; *ROSA26*^*mT/mG*^ mutants showed fewer GFP^+^ cNCCs in the PA1 (yellow arrow), PA2 and migratory stream relative to control embryos at E10.5. (C) Apoptosis was examined at E10.5 by TUNEL. Increased TUNEL positive cells were observed *Fgfr1*^*cKO/cKO*^; *Fgfr2*^*cKO/cKO*^ mutants in the lateral nasal process (LNP) compared to controls. Fewer TUNEL positive cells were observed in the medial nasal process (MNP). (D) Quantitation of TUNEL positive foci across different mutant genotypes show a 40-fold increase in cell death in *Fgfr1/2* double mutants in the LNP (“*” inside the bar graph represents significance level compared to control) at E10.5. (E) Inferior view (mandibles removed) of alcian blue/ alizarin red stained mouse skulls in *Fgfr1*^*cKO/cKO*^; *Fgfr2*^*cKO*/+^; *Bim*^+/+^ and *Fgfr1*^*cKO/cKO*^; *Fgfr2*^*cKO*/+^; *Bim*^+/−^ embryos showed partial rescue of medial structures, the nasal cartilage (NC) and palatine (PL) process, premaxilla (PMX), maxilla (MX), basisphenoid (BS) and palatal shelves (PS) in *Bim*^−/+^ mutants. Black bar (-) measures midline separation. Average midline separation (mm) is reduced by 2-fold (*N*=4, P>0.0109) in *Fgfr1*^*cKO/cKO*^; *Fgfr2*^*cKO*/+^; *Bim^+/−^* as represented in the graph. (F) Frontal views of control *Fgfr1*^*cKO*/+^; *Fgfr2*^*cKO*/+^; *Bim*^+/+^, *Fgfr1*^*cKO/cKO*^; *Fgfr2*^*cKO/cKO*^; *Bim*^+/+^, and *Fgfr1*^*cKO/cKO*^; *Fgfr2*^*cKO/cKO*^; *Bim*^+/−^ embryos at E17.5 showing partial phenotypic rescue. Black bar (**−**) indicates intercanthal distance.

A reduction in NCC numbers in the midface might occur due to reduced proliferation, increased cell death, or both. Cell proliferation remained unaffected (Extended Data Fig. 3), however, we observed enhanced cell death in *Fgfr1*^*cKO/cKO*^; *Fgfr2*^*cKO/cKO*^; *ROSA26*^*mT/mG*^ E10.5 mutants in the craniofacial mesenchyme, notably in the Lateral Nasal Process (LNP) relative to the Medial Nasal Process (MNP; Fig. 1C, D). We next asked to what extent this observation could explain the overall morphological defects. The BH3-only protein BIM plays a critical role in initiating apoptotic pathway in multiple cell types by binding and repressing the function of several pro-survival BCL-2 family members ^12–14^, and is involved in craniofacial development ^15^. Consistent with a role for cell survival in the etiology of craniofacial defects, we found that *Fgfr1*^*cKO/cKO*^; *Fgfr2*^*cKO*/+^; *Bim*^+/−^ E17.5 mutants exhibited a 55% reduction in midline separation along with a partial rescue of medial skeletal structures, including the anterior nasal cartilage, palatine process, premaxilla, primary and secondary palate, pterygoid process and basisphenoid bone, compared to control embryos (Fig. 1E). The reduction in *Bim* levels significantly rescued defects particularly at the level of the nasal mesenchyme and the midface (Fig. 1F and Supplementary Information Fig. 1E). BIM is a known target of phosphorylation by several MAP kinases, particularly ERK1/2 which targets it for ubiquitination and proteasomal degradation ^16^. JNK and PI3K/AKT activation are also known to affect BIM levels ^17^. Cell survival through BIM may therefore be regulated by FGF since several of these signaling pathways are engaged by FGFR1 and FGFR2 ^5^. Taken together, these results underscore the importance of cell death in the conditional double null mutant phenotype.

## Combined *Fgfr1/2* signaling mutations do not recapitulate the null phenotypes

To interrogate signaling mechanisms *in vivo*, we generated an allelic series of knock-in point mutations at the *Fgfr2* locus preventing binding of effectors to the receptor (Fig. 2A-B; Extended Data Fig. 4), similar to previous *Fgfr1* mutations ^7^. The *Fgfr2*^*F*^, *Fgfr2*^*C*^, and *Fgfr2*^*PG*^ mutations were designed to disrupt binding of FRS2, CRK-L and PLCγ/GRB14, respectively. We also generated compound *Fgfr2*^*CPG*^ and *Fgfr2*^*FCPG*^ signaling mutants by combining multiple signaling mutations. Co-immunoprecipitation and western blot analysis showed that each signaling mutation disrupted effector binding (Fig. 2B). *Fgfr2* signaling mutant alleles were evaluated for their ability to partially or completely recapitulate the *Fgfr2*^−/−^ E10.5 placenta and limb phenotype ^18–20^. Surprisingly, all signaling allele mutants were at least partially viable and fertile as homozygotes (Supplementary Information Table S1 and Supplementary Information Fig. S2). *Fgfr2*^*F/F*^ and *Fgfr2*^*FCPG/FCPG*^ homozygous mutants also exhibited defects in lacrimal gland branching morphogenesis, a process regulated by FGF signaling ^21,22^ (Extended Data Fig. 5). IRS2 has been found to bind FGFR2 ^23^, but no genetic interaction was observed between *Irs2*^−/−^ and *Fgfr2*^*FCPG/FCPG*^ mutants.

**Figure 2:**
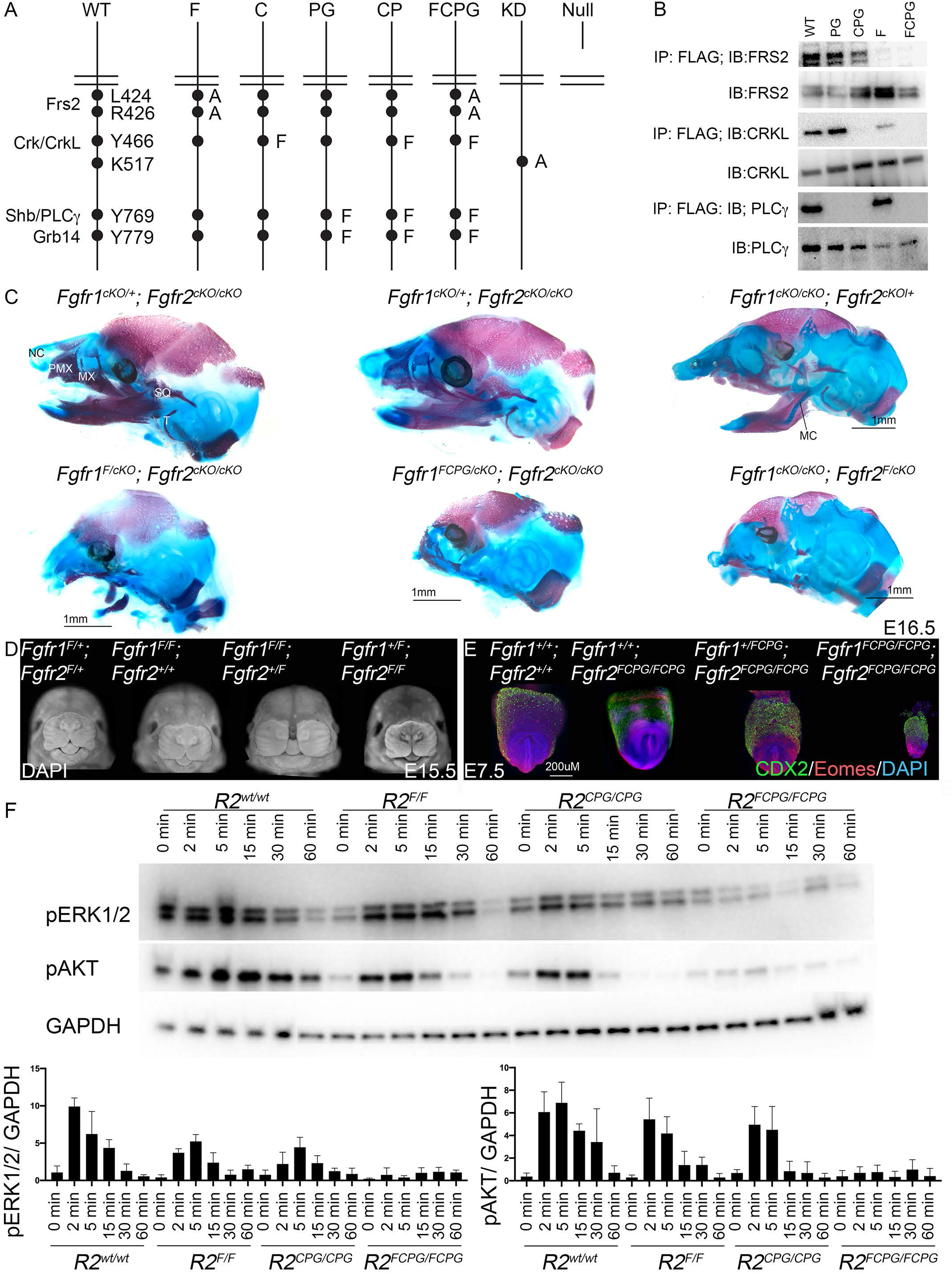
Coordinate roles *Fgfr1* and *Fgfr2* signaling mutation in development. (A) Schematic representation of the *Fgfr2* allelic series. Critical effectors that bind to FGFR2 are listed on the left of the *Fgfr2*^*WT*^ allele (WT). Critical residues for this binding are annotated on the right. Amino acid substitutions for each allele are provided to the right of all mutant alleles generated. (B) Co-immunoprecipitation experiments confirmed the ability of F, C and PG mutations to disrupt FRS2, CRKL and PLCγ binding to FGFR2, respectively. Constructs expressing *Fgfr2^WT^, Fgfr2^PG^, Fgfr2^CPG^, Fgfr2^F^* and *Fgfr2*^*FCPG*^-triple Flag-tag cDNA were overexpressed in 3T3 cells. FLAG pull-downs were then immunoblotted to show interactions of FRS2, CRKL and PLCγ to FGFR2^WT^ and mutant receptors. (C) Sagittal view of E16.5 alcian blue/ alizarin red stained skulls from control littermate (top) and *Fgfr1* or *Fgfr2* signaling (bottom) mutants. In *Fgfr1^F/cKO^; Fgfr2^cKO/cKO^* mutants, the frontal bone, nasal cartilage (NC), squamosal bone (SQ), tympanic bulla (T), maxilla (MX) and mandible (MD) were affected. *Fgfr1^FCPG/cKO^; Fgfr2^cKO/cKO^* mutants showed a more severe defect. *Fgfr2^F^* signaling mutants (*Fgfr1^cKO/cKO^; Fgfr2^F/cKO^*) showed defects in mandible along with reduced ossification of the premaxilla (PMX) and maxilla and loss of the squamosal bone and tympanic bulla. (D) DAPI stained frontal facial view for *Fgfr1^F^* and *Fgfr2^F^* mutant embryos at E15.5. *Fgfr1^F/F^; Fgfr2^+/F^* embryos showed a severe facial cleft compared to other genotypes. (E) CDX2 and EOMES expression in *Fgfr1^FCPG^* and *Fgfr2*^*FCPG*^ compound signaling mutants at E7.5. In contrast to *Fgfr1^−/−^; Fgfr2^−/−^* which fail to implant on the 129S4 co-isogenic background, we could recover *Fgfr1^FCPG/FCPG^; Fgfr2^FCPG/FCPG^* compound mutants at E7.5 but not at E10.5. *Fgfr1^FCPG/FCPG^; Fgfr2^FCPG/FCPG^* mutants were growth-retarded but expressed mesodermal EOMES and the trophectoderm marker CDX2. (F) iFNP cells (*Fgfr1^CRISPR-KO^*) derived from the indicated genotypes, *Fgfr2^wt/wt^* (*R2^wt/w^*), *Fgfr2^F/F^* (*R2^F/F^*), *Fgfr2^CPG/CPG^* (*R2^CPG/CPG^*) and *Fgfr2^FCPG/FCPG^* (*R2^FCPG/FCPG^*), were stimulated with FGF1 for the indicated times (0-60 mins) to analyze pERK1/2 and pAKT activation. Quantification of pathway activation for pERK1/2 and pAKT, normalized to GAPDH, is reported as mean ± standard deviation with a minimum of three independent biological replicates.

Because *Fgfr1* and *Fgfr2* are co-expressed in cNCCs, we reasoned that discrete functions of signaling pathways downstream of one receptor could be masked by the presence of the other, wild-type receptor. To test this hypothesis, we analyzed compound conditional hemizygous *Fgfr1^F^*, *Fgfr1^FCPG^* and *Fgfr2^F^* mutations over the *Fgfr1^cKO^* and *Fgfr2^cKO^* conditional null alleles. *Fgfr1^F/cKO^; Fgfr2^cKO/cKO^* and *Fgfr1^FCPG/cKO^; Fgfr2^cKO/cKO^* conditional mutants developed severe agenesis of midface structures, nasal cartilage, and mandible but the phenotype was less severe than in *Fgfr1*^*cKO/cKO*^; *Fgfr2*^*cKO/cKO*^ mutants. The nasal cartilage and mandible defects were further exacerbated in *Fgfr1^FCPG/cKO^; Fgfr2^cKO/cKO^* mutants (Fig. 2C, Extended Data Fig. 6A, red arrows, and Supplementary Information Table S2). *Fgfr1^cKO/cKO^; Fgfr2^F/cKO^* embryos developed more severe midline fusion and mandible defects than *Fgfr1*^*cKO/cKO*^; *Fgfr2*^*cKO*/+^ controls. Similar to *Fgfr1*^*cKO/cKO*^; *Fgfr2*^*cKO/cKO*^ mutants, morphological defects in *Fgfr1^F/cKO^; Fgfr2^cKO/cKO^* and *Fgfr1^FCPG/cKO^; Fgfr2^cKO/cKO^* embryos arose as early as E10.5 and were accompanied by cell death in the LNP (Extended Data Fig. 6B-E).

We next examined the importance of coordinate engagement of signaling pathways by both receptors upon ligand activation. *Fgfr1^C/C^; Fgfr2^C/C^* and *Fgfr1^CPG/CPG^; Fgfr2^CPG/CPG^* double mutants were recovered in normal numbers, fertile, and did not exhibit a craniofacial phenotype. Compound *Fgfr1^F/F^; Fgfr2^+/F^* mutants displayed hypoplastic nasal prominences and midline fusion defects at E15.5 (Fig. 2D) with no defect in mandibular development (Extended Data Fig. 6F), but *Fgfr1^F/F^*; *Fgfr2^F/F^* mutant embryos were not recovered at E10.5. Similar hypoplastic nasal and mandibular prominence defects were observed in *Fgfr1^FCPG/FCPG^; Fgfr2^+/FCPG^* compound mutants (Extended Data Fig. 6G). Interestingly, *Fgfr1^FCPG/FCPG^; Fgfr2^FCPG/FCPG^* mutants survived to E7.5 and still formed mesoderm, as evidenced by *Eomes* staining (Fig. 2E), in contrast to *Fgfr1^−/−^; Fgfr2^−/−^* double mutants which fail at implantation on the same genetic background ^24^. These results indicate that signaling mutations in both *Fgfr1* and *Fgfr2* interact genetically during development, but that the combination of the most severe signaling mutations fails to recapitulate the double null mutant phenotype.

## *Fgfr1/2* signaling mutations abrogate signal transduction cascades

FGFs activate numerous signaling pathways upon ligand stimulation ^5^. To evaluate intracellular pathway activation downstream of wild-type FGFR2, FGFR2^F^, FGFR2^CPG^, and FGFR2^FCPG^, we generated immortalized E10.5 frontonasal prominence cell lines (iFNPs) by crossing to *Ink4a/Arf* mutants ^25^. These cells express predominantly *Fgfr1* and to a lesser extent *Fgfr2*, but no *Fgfr3* or *Fgfr4*. We then eliminated FGFR1 expression by CRISPR/Cas9 mutagenesis leaving FGFR2 as the sole receptor, and interrogated activation of six pivotal FGF-engaged pathways ^5^ by FGF1, which gave more robust responses than FGF8b (Extended Data Fig. 7). For ERK1/2, but also for PI3K/AKT and PLCγ, robust activation seen in *wt*-iFNPs was diminished in both *Fgfr2^F/F^* or *Fgfr2^CPG/CPG^*-iFNPs, but only eliminated in *Fgfr2^FCPG/FCPG^* mutant cells (Fig. 2F and Extended Data Fig. 8). Our data thus indicates that both FRS2 and CRKL/PLCγ binding is necessary for activation of these pathways, as for *Fgfr1* ^7^. P38, STAT3 and JNK activation were also abrogated in *Fgfr2^FCPG/FCPG^*-iFNPs, with variable effects in *Fgfr2^F/F^*-iFNPs and *Fgfr2^CPG/CPG^*-iFNPs (Extended Data Fig. 8). Last, we revisited cell signaling pathways in *Fgfr1^FCPG/FCPG^* mutants ^7^. We used CRISPR/Cas9 to create *Fgfr1^FCPG/FCPG^*; *Fgfr2^CRISPR-KO^*-iFNP cells, and found that both ERK1/2 and pAKT activation were eliminated in these cells (Extended Data Fig. 8F-G). Together, these results indicate that the most severe signaling mutation combinations in *Fgfr1* and *Fgfr2* effectively abrogate classic signal transduction pathways for each receptor.

## *Fgfr2* function requires its kinase activity

The lack of more severe phenotypes in *Fgfr2* signaling mutants raised the possibility that FGFR2 acts independent of kinase activity. To address this question, we generated a K517A kinase dead (KD) mutation in the ATP binding site at the *Fgfr2* locus ^26,27^ (Fig. 2A and Fig. 4A, C). *Fgfr2^KD/+^* heterozygous embryos (27/60) showed no obvious defects at E10.5. *Fgfr2^KD/+^* heterozygotes appeared normal during later developmental stages up to P0, but fewer than expected (24/78) were recovered after weaning. They exhibited no limb or craniofacial abnormalities, but significantly more severe semi-dominant defects in lacrimal gland development than *Fgfr2^F/F^* and *Fgfr2^FCPG/FCPG^* mutants by P15 (Extended Data Fig. 5A-C).

No *Fgfr2^KD/KD^* homozygotes were recovered at birth (Supplementary Information Table S1). Morphological examination of *Fgfr2^KD/KD^* embryos at E10.5 showed characteristic *Fgfr2^−/−^* phenotypes, with absence of limb buds, incomplete connection of the allantois to the ectoplacental cone, and dilated pericardium (Fig. 3A), suggesting that FGFR2 broadly operates in a kinase-dependent fashion. Moreover, since the *Fgfr2b* constitutive mutation leads to similar limb phenotypes but no placental insufficiency ^11^, and phenotypes in both tissues are observed in *Fgfr2^−/−^* or *Fgfr2^KD/KD^* mutants, FGFR2 activity must be kinase-dependent in both mesenchymal and epithelial contexts. However, *Fgfr2^KD/KD^* embryos exhibited additional phenotypes including severe posterior truncations and craniofacial defects (Fig. 3A). We tested for complementation between *Fgfr2^KD^* and *Fgfr2^−^* alleles to evaluate phenotypic differences. *Fgfr2^KD/−^* embryos showed absence of limb buds and defects in the chorio-allantoic junction, along with dilated pericardium, similar to *Fgfr2^−/−^* embryos, and posterior truncations defects similar to *Fgfr2^KD/KD^* embryos (Fig. 3A). In contrast, defects in the forebrain, medial and lateral nasal prominences, and maxillary and mandibular prominences appeared less severe than in *Fgfr2^KD/KD^* mutants (Fig. 3B-D).

**Figure 3:**
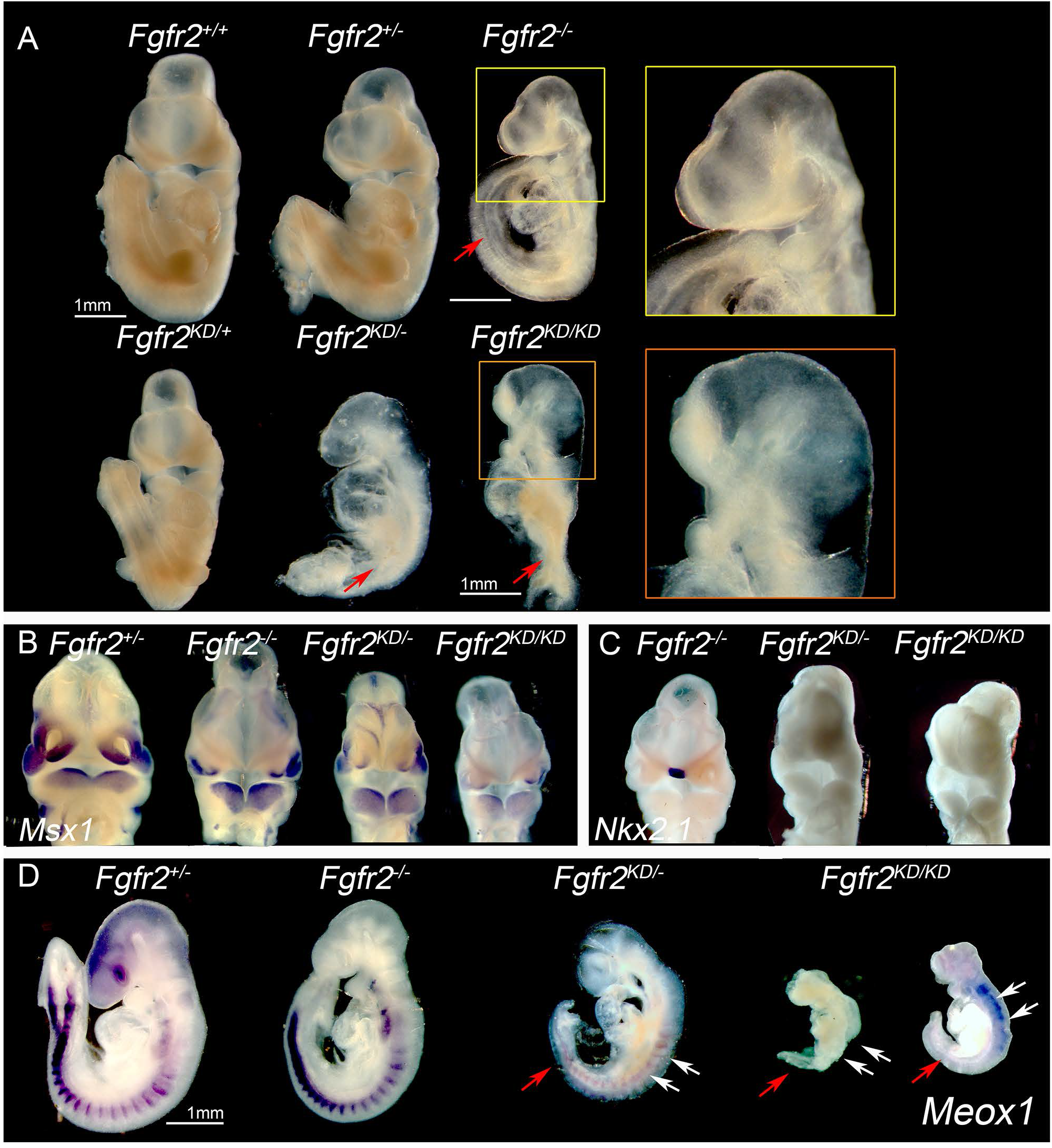
A kinase dead mutation in *Fgfr2* recapitulates multiple aspects of the *Fgfr2*^−/−^ phenotype. (A) Morphological examination of *Fgfr2* kinase dead phenotype at E10.5. *Fgfr2^KD/−^* and *Fgfr2^KD/KD^* mutants had no limb buds (red arrows), and had defects in the allantois which was loosely or incompletely held by the chorion to the ectoplacental cone, along with a dilated pericardium similar to *Fgfr2^−/−^* mutants. Both *Fgfr2^KD/−^* and *Fgfr2^KD/KD^* mutants displayed severe posterior truncations and *Fgfr2^KD/KD^* mutants showed a more severe phenotype. *Fgfr2^KD/KD^* mutants also displayed craniofacial defects with poorly developed facial prominences compared to *Fgfr2^KD/−^* mutants. (B) Wholemount mRNA *in situ* hybridization for cNCC mesenchyme marker *Msx1* at E10.5. *Msx1* was expressed in the facial prominences (LNP, MNP and mandibular prominences) in *Fgfr2*^−/−^ embryos at similar levels compared to control *Fgfr2^+/−^* embryo. *Msx1* expression was also observed in *Fgfr2^KD/−^* mutants but was reduced in *Fgfr2^KD/KD^* mutants. (C) Wholemount mRNA in situ hybridization for midline marker *Nkx2.1*. *Nkx2.1* was expressed in the midline floor plate of *Fgfr2^−/−^* mutants, but not detected in *Fgfr2^KD/−^* and *Fgfr2^KD/KD^* mutants. (D) *Meox1* mRNA expression in somites of E10.5 *Fgfr2^+/−^* embryos was comparable to *Fgfr2^−/−^* mutants, but severely reduced in both *Fgfr2^KD/−^* and *Fgfr2^KD/KD^* mutants (white arrows) along with defects in axis elongation and posterior somite formation (red arrow).

FGFRs can form heterodimers in culture ^27,28^, although these have never been demonstrated *in vivo* in the absence of over-expression. The semi-dominant effects observed in *Fgfr2^KD/+^* heterozygous mutants affecting postnatal lacrimal gland development, and the craniofacial and mesoderm defects observed in *Fgfr2^KD/KD^* embryos suggest that the *Fgfr2^KD^* allele not only inactivates *Fgfr2* but also suppresses *Fgfr1* activity through FGFR2^KD^:FGFR1 heterodimers, wherever they are co-expressed. This would explain why *Fgfr2^KD^* mutants still generally resemble *Fgfr2^−/−^* mutants which are lethal at a similar stage, rather than *Fgfr1^−/−^* mutants^7,18,24^. While signaling might occur through heterodimers, this mechanism cannot fully account for the discrepancy between FCPG and null phenotypes for either receptor, as *Fgfr1^FCPG/FCPG^; Fgfr2^FCPG/FCPG^* double mutants develop until E7.5 with a significant degree of mesoderm formation (Fig. 2E) whereas *Fgfr1; Fgfr2* double null mutants have a more severe defect at implantation ^18,24^. Alternatively, the FGFR2^KD^ receptor could act as a dominant negative by titrating/sequestering ligand away from functional receptors, if it is distributed differently in the cell, but this is unlikely as *Fgfr2^FCPG/+^* mutants do not show dominant negative effects and both knock-in alleles should be expressed at similar levels.

## Cell matrix and cell adhesion properties are retained beyond canonical signaling

Since signaling mutant cells showed near complete inactivation of classic RTK signaling activities, but the corresponding mutant mice failed to recapitulate the null mutant phenotype, we reasoned that some function engaged by FGF signaling must be retained in the most severe *FCPG* mutants. Previous lines of evidence have implicated FGF signaling in the control of cell-matrix ^29^ or cell-cell adhesion ^24,30,31^. We examined *Fgfr1*; *Fgfr2* dependent cell spreading/migration on extracellular matrix across a wound, using primary *Fgfr1^cKO/+^; Fgfr2^cKO/+^*, *Fgfr1^FCPG/+^; Fgfr2^cKO/+^*, or *Fgfr1*^*cKO/cKO*^; *Fgfr2*^*cKO/cKO*^ FNP cells. Spreading of control cells over the wound over twelve hours was comparable upon FGF, PDGF, and serum stimulated conditions. Interestingly, *Fgfr1^FCPG/cKO^; Fgfr2^cKO/cKO^* cells showed comparable spreading capacities in all three stimulation conditions. However, *Fgfr1*^*cKO/cKO*^; *Fgfr2*^*cKO/cKO*^ double mutant cells failed to spread into the wound area in response to FGF, while responses to PDGF and serum were normal (Fig. 4A).

**Figure 4:**
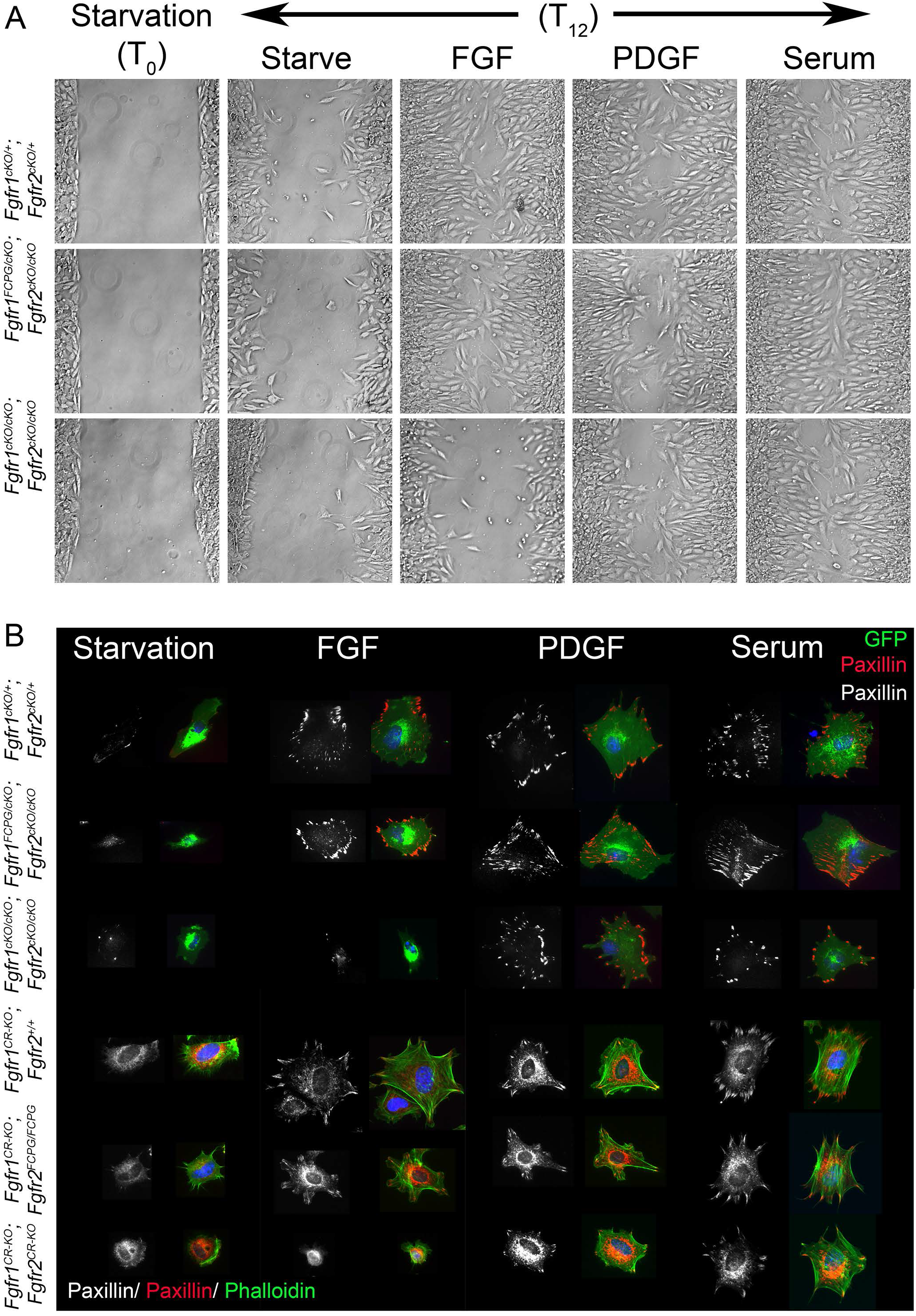
Cell matrix adhesion properties are retained in FCPG mutants. (A) Scratch/wound healing assay was used to study cell spreading. Control (*Fgfr1^cKO/+^; Fgfr2^cKO/+^*) and *Fgfr1^FCPG/cKO^; Fgfr2^cKO/cKO^* primary FNP cells showed active spreading over12 hours (T12) in response to FGF1, PDGF-A, or serum. *Fgfr1*^*cKO/cKO*^; *Fgfr2*^*cKO/cKO*^ mutant cells showed limited spreading upon FGF stimulation but normal spreading in response to PDGF or serum. (B) Focal-adhesion formation was assayed by Paxillin immunostaining in *Fgfr1* and *Fgfr2* signaling mutant primary FNP cells. GFP^+^ *Fgfr1^cKO/+^; Fgfr2^cKO/+^; ROSA26^mT/mG^* control, *Fgfr1^FCPG/cKO^; Fgfr2^cKO/+^; ROSA26^mT/mG^,* and *Fgfr1^cKO/cKO^; Fgfr2^cKO/cKO^; ROSA26^mT/mG^* mutant cells were treated with FGF1, PDGFA or serum for 3 hours before analyzing Paxillin localization to focal adhesions. In control cells and *Fgfr1^FCPG/cKO^; Fgfr2^cKO/+^*, we detected multiple Paxillin^+^ foci upon FGF1, PDGFA, or serum stimulation. *Fgfr1^cKO/cKO^; Fgfr2^cKO/cKO^; ROSA26^mT/mG^* mutant FNP cells failed to form focal adhesions in response to FGF, in contrast to PDGF or serum (quantified in Extended Data Fig. 9A). Cell spreading properties were also analyzed for *Fgfr2^WT/WT^* (*Fgfr1CRISPR-KO*; *Fgfr2WT/WT*), *Fgfr2FCPG/FCPG* (*Fgfr1CRISPR-KO*; *Fgfr2FCPG/FCPG*), and *Fgfr2−/−* (*Fgfr1CRISPR-KO*; *Fgfr2 ^CRISPR-KO^*)-iFNP cells in which *Fgfr1* was disrupted. Addition of FGF1, PDGFA or serum resulted in robust cell spreading and formation of Paxillin^+^ focal adhesion in *Fgfr1^CRISPR-KO^*; *Fgfr2^FCPG/FCPG^* and *Fgfr1^CRISPR-KO^*; *Fgfr2^+/+^*-iFNP cells. *Fgfr2^−/−^* (*Fgfr1^CRISPR-KO^*; *Fgfr2 ^CRISPR-KO^*)-iFNP cells showed severe defects in focal adhesion formation upon FGF1 treatment, but was unaffected in response to PDGFA or serum (quantified in Extended Data Fig. 9E).

We next investigated focal adhesion formation and found that GFP^+^ *Fgfr1^cKO/+^; Fgfr2^cKO/+^; ROSA26^mT/mG^* control cells and *Fgfr1^FCPG/cKO^; Fgfr2^cKO/cKO^; ROSA26^mT/mG^* mutant cells formed numerous Paxillin enriched focal adhesions in response to FGF, PDGF or serum. Double null mutant cells however failed to form any Paxillin^+^ foci upon FGF treatment, but still responded normally to PDGF and serum as expected (Fig. 4B and Extended Data Fig. 9A). Western blot analysis showed reduced pFAK and total Paxillin levels in *Fgfr1*^*cKO/cKO*^; *Fgfr2*^*cKO/cKO*^ cells compared to *Fgfr1^cKO/+^; Fgfr2^cKO/+^* or *Fgfr1^FCPG/cKO^; Fgfr2^cKO/cKO^* cells, and FAK phosphorylation depended on FGF (Extended Data Fig. 9B-D). Likewise, we analyzed *Fgfr1^CRISPR-KO^; Fgfr2^+/+^* and *Fgfr1^CRISPR-KO^; Fgfr2^FCPG/FCPG^*-iFNP cells and found that they spread and formed Paxillin^+^ focal adhesions in response to serum, PDGF, or FGF, in contrast to *Fgfr1^CRISPR-KO^; Fgfr2^−/−^*-iFNP cells (Fig. 4B and Extended Data Fig. 9E). These observations indicate that although FGFR1^FCPG^ and FGFR2^FCPG^ lose most FGF-dependent intracellular kinase signaling outputs, they still retain functions pertaining to cell-matrix interactions and that cell spreading and stabilization of cell-matrix interactions are actively governed by FGF signaling to its receptors.

We also explored if *Fgfr1^FCPG^* cells established stable cell-cell contacts comparable to control cells. We found extensive adherens junctions among *Fgfr1^cKO/+^; Fgfr2^cKO/+^; ROSA26^mT/mG^* control cells and *Fgfr1^FCPG/cKO^; Fgfr2^cKO/cKO^; ROSA26^mT/mG^* cells, marked by localized β-catenin along cell boundaries (Fig. 5A-B and Extended Data Fig. 9F). In contrast, double null mutant cells showed far fewer cell-cell contacts with no localized β-catenin accumulation, suggesting that contacts are either unstable or do not mature (Fig. 5B). Last, we examined cell-cell contacts *in vivo*, in the E11.5 LNP. GFP^+^ cNCCs in the mesenchyme exhibited extensive cell-cell contacts within the LNP in both control and *Fgfr1^FCPG/cKO^; Fgfr2^cKO/cKO^; ROSA26^mT/mG^* embryos. Strikingly, GFP^+^ cNCCs in the double null mutant LNP were mostly isolated and interspersed, whereas cell contacts in the MNP remained unaffected (Fig. 5C). Taken together, these results indicate that the most severe signaling mutations in *Fgfr1* and *Fgfr2* still retain cell-matrix and cell-cell interactions otherwise lost in the nulls, while abrogating classic signal transduction pathways.

**Figure 5:**
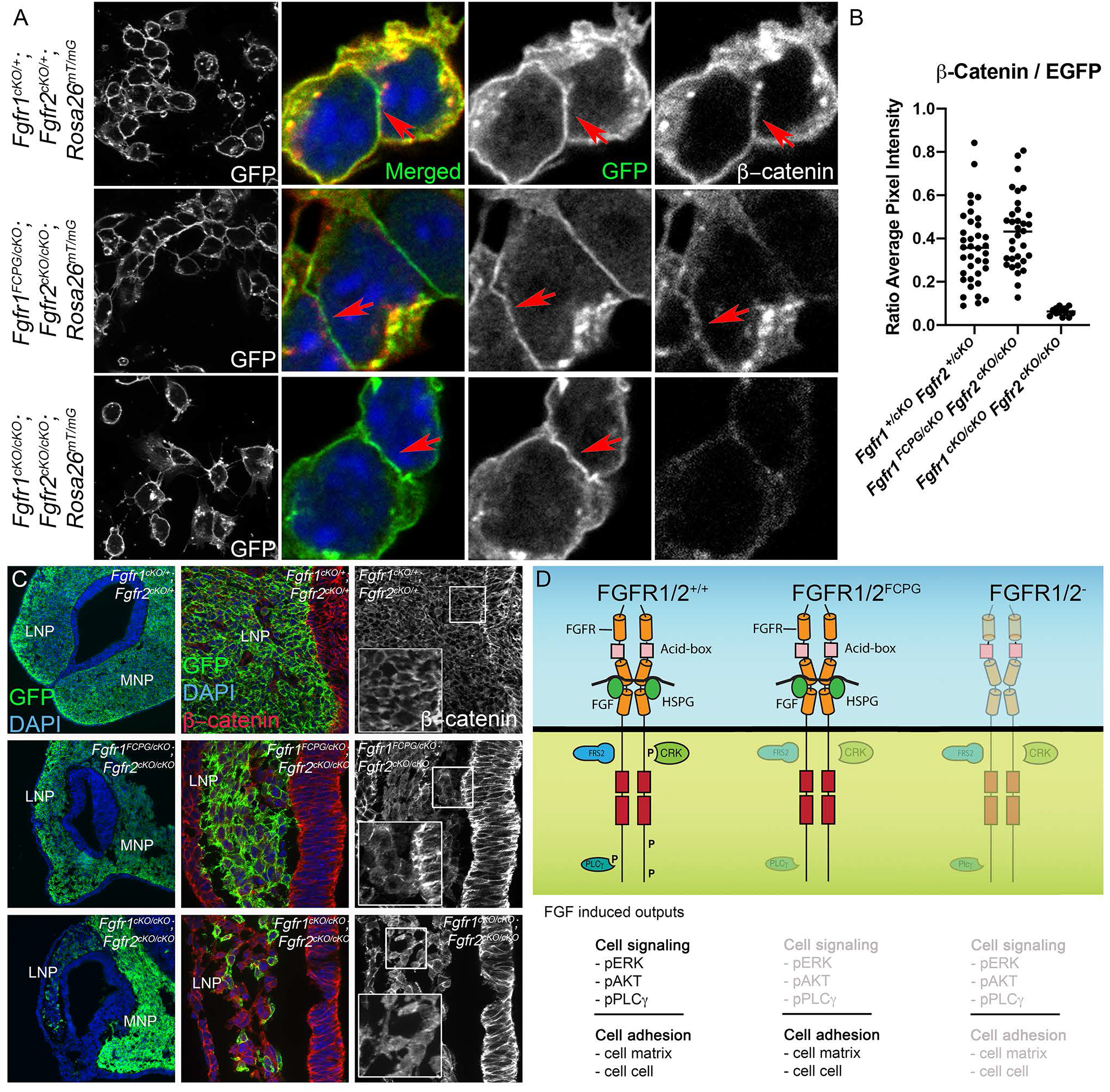
Cell-cell adhesion is retained in *Fgfr* signaling mutants. (A) GFP^+^ primary FNP cells from control *Fgfr1^cKO/+^; Fgfr2^cKO/+^; ROSA26^mT/mG^* and *Fgfr1^FCPG/cKO^; Fgfr2^cKO/cKO^; ROSA26^mT/mG^* embryos formed extensive cell-cell contacts, in contrast to *Fgfr1^cKO/cKO^; Fgfr2^cKO/cKO^; ROSA26^mT/mG^* mutants. Quantification is provided in Extended Data Fig. 9F. β-catenin was localized along the cell contact boundaries in GFP^+^ FNP cells from control and *Fgfr1^FCPG/cKO^; Fgfr2^cKO/cKO^; ROSA26^mT/mG^* embryos in contrast to *Fgfr1^cKO/cKO^; Fgfr2^cKO/cKO^; ROSA26^mT/mG^* mutants. (B) β-catenin immunofluorescence was quantified and represented as mean for each genotype. (C) Extensive cell-cell contacts with β-catenin localization (insets) were observed *in vivo* in control *Fgfr1^cKO/+^; Fgfr2^cKO/+^; ROSA26^mT/mG^* and *Fgfr1^FCPG/cKO^; Fgfr2^cKO/cKO^; ROSA26^mT/mG^* cellsin both developing LNP and MNP at E11.5. In contrast, *Fgfr1^cKO/cKO^; Fgfr2^cKO/cKO^; ROSA26^mT/mG^* double null mutants GFP^+^ cells remained sparse in the LNP and showed no localized β-catenin. (D) Model of FGF-mediated cell signaling pathways. In wild-type (WT) cells, activation by FGFs engages a canonical RTK signal transduction pathway, leading for the activation of ERK1/2, PI3K/AKT, PLCγ, and additional pathways. In addition, FGFs activate non-canonically both cell-matrix as well as cell-cell adhesion, in a kinase-dependent manner, possibly facilitated through interactions of the FGF receptors through their extracellular domain with cell adhesion receptors. *Fgfr1^FCPG^* or *Fgfr2*^*FCPG*^ mutant cells fail to activate a classical RTK signal transduction pathway (light grey) but can still promote cell adhesion (black), as their kinase activity has not been disrupted. In null mutant cells, neither FGF-induced cell signaling nor cell adhesion are observed (light grey), since the receptors are not expressed.

## Discussion

FGFs are best known for activating signal transduction cascades, most prominently ERK1/2 ^5,6^, Surprisingly, we find that mutations in *Fgfr1* and *Fgfr2* which broadly eliminate classic RTK signaling outputs fail to recapitulate the *Fgfr1^−/−^* or *Fgfr2^−/−^* phenotypes. The near complete abrogation of multiple signal transduction outputs in the most severe *Fgfr1* and *Fgfr2* signaling alleles indicates that we have interrogated relevant cell signaling pathways and implies the existence of non-canonical functions not impacted by our signaling mutations. Importantly, while the *FCPG* mutations disrupt the ability of intracellular effectors to engage classical RTK activity, they do not abrogate the receptor’s kinase activity, suggesting that FGFR1^FCPG^ and FGFR2^FCPG^ receptors may be able to phosphorylate other targets.

A wide body of literature has implicated FGFRs in cell adhesion through interactions of the extracellular domain, which remains untouched in any of our signaling mutations, with cell adhesion molecules. FGF ligand binding to FGFRs is known to involve a third party, heparan sulfate proteoglycans (HSPGs)^32,33, 34^, which in turn interact with integrins, regulating cell-matrix adhesion ^35–37^. Consistent with a critical role for FGFRs, we found that cell-matrix adhesion was unperturbed in cells carrying the most severe signaling mutations, but lost in the absence of FGFRs following FGF stimulation. Because FGF ligand still promotes cell-matrix adhesion in cells derived from *Fgfr1/2^FCPG^* mutants, the activity of cell adhesion receptors must be enhanced independent of traditional signaling outputs, either directly through their interaction with FGFRs or indirectly by an unknown molecule/kinase. Defects in FGF-dependent adhesion could also result in the induction of anoikis ^38^, a process previously shown to be regulated by *Bim* ^39^, potentially linking our observed adhesion defects and increase in cell death in the LNP. Last, focal adhesion assembly and phosphorylation defects have also been observed in *Fgfr1^−/−^; Fgfr2^−/−^* keratinocytes ^29^, suggesting that engagement of FGF signaling has a broad function in regulating cell adhesion in both mesenchymal and epithelial contexts. Other receptor tyrosine kinases, like PDGFRs, have been found to bind integrins ^40^ and PDGFR signaling mutations have also failed to recapitulate null mutant phenotypes ^41,42^.

In addition, both FGFR1 and FGFR2 are known to interact through their extracellular domain acid box with various cell adhesion molecules such as cadherins ^43^, critical players in cNCC migration ^44^. Consistent with a role for FGFRs in mediating cell-cell adhesion through cadherins, we observed that cells derived from the most severe cell signaling mutants made strong β-catenin cell contacts, in contrast to double null mutant cells. We also observed fewer contacts *in vivo*, where double null mutant cells showed very limited interactions in the LNP. FGFR1/2 association with cadherins may thus have a dual role, one in promoting cell motility through FGF-induced ERK1/2 signaling ^43^, and an opposite one in cell adhesion, as ERK1/2 signaling is abrogated in *Fgfr1^FCPG^* and *Fgfr2*^*FCPG*^ mutants. FGF signaling regulates E-cadherin localization, and in the absence of *Fgfr1*, E-cadherin polarization is affected in mural trophectoderm ^24^, *Drosophila* mesoderm ^31^, and zebrafish cardiomyocytes ^30^. FGF signaling also regulates cadherin switching during epithelial to mesenchymal transition, including neural crest cell delamination ^45–47^. Taken together, our results indicate that FGFRs regulate cell adhesion, and possibly other processes, beyond their classic signaling cascades (Fig. 5D). Additional genetic, biochemical and cell biological studies may identify further non-canonical roles for these receptors beyond their traditional activities in signal transduction.

## Acknowledgments

We thank Jia Li and Chantel Dixon for technical assistance; Kevin Kelley for stable tissue culture facilities; Elaine Fuchs for helpful insights into cell adhesion mechanisms; Jerry Chipuk for conversations about cell death; and our laboratory colleagues, Stu Aaronson, Rob Krauss, and Sergei Sokol for critical comments on the manuscript. We thank the NYU School of Dentistry Micro-CT Core and the Mt. Sinai Flow Cytometry and Microscopy facilities for assistance and advice. C.J.D was supported by F32 DE026678 from NIH/NIDCR. This work was supported in part by the Tisch Cancer Institute at Mount Sinai (P30 CA196521 Cancer Center Support Grant, for access to Mt. Sinai cores) and by grant RO1 DE022778 from NIH/NIDCR to P.S.

## Figures, Extended Figures and Supplementary Information

**Extended Data Fig. 1:**
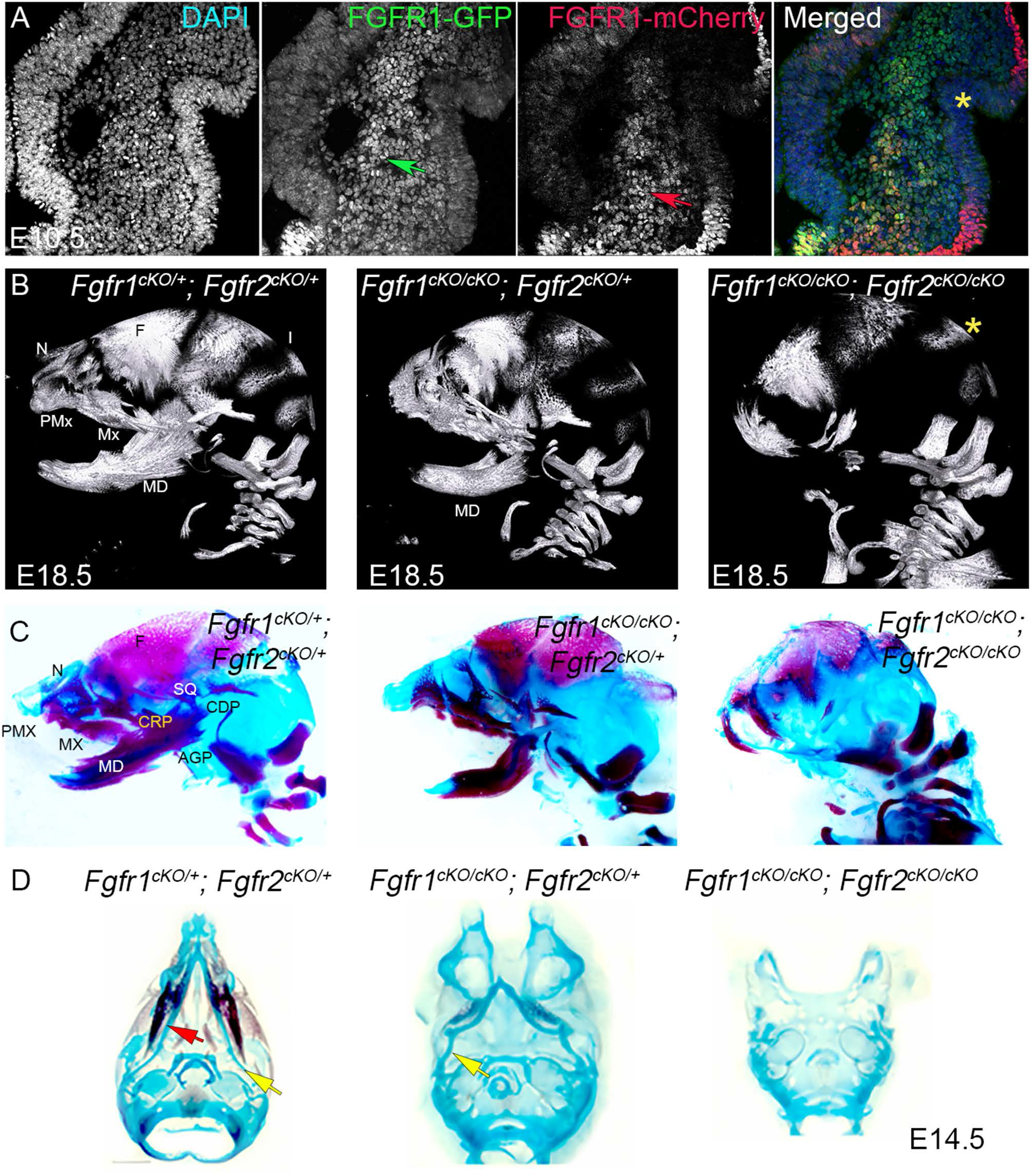
Defects in craniofacial morphogenesis in *Fgfr1*^*cKO/cKO*^; *Fgfr2*^*cKO/cKO*^ double mutants. (A) Spatial domain of FGFR1 and FGFR2 expression in facial prominences at E10.5. GFP and mCherry immunohistochemistry were used to detect expression from *Fgfr1-GFP* and *Fgfr2-mCherry* reporter alleles. GFP expression was primarily restricted to the mesenchyme (green arrow). Although mCherry expression was restricted to the epithelium, many cells in the mesenchyme also express mCherry (red arrow). mCherry expression is downregulated in the epithelium lining the nasal pit (yellow asterisk). FGFR1 and FGFR2 co-expression in the craniofacial skeleton is also observed E17.5 (Supplementary Information Fig. S1a). (B) Micro-CT analysis of ossified structures in the head at E18.5. An overall reduction or complete loss of ossification of cNCC-derived facial bones in *Fgfr1/2* conditional double mutant is seen compared to the control. Frontal (F) and nasal bones (N), pre-maxilla (PMX) and maxilla (MX) were severely affected. Mesoderm derived bones including parietal and interparietal(I) bones remained unaffected (yellow asterisk). *Fgfr1*^*cKO/cKO*^; *Fgfr2*^*cKO*/+^ mutants showed an overall reduction in the size of the mandible (MD), with the proximal structures most severely affected. A more detailed morphometry analysis regarding changes in skull length, intercanthal distance, length of frontal bone and length of nasal cartilage at E18.5 for different genotypes is provided in Supplementary Information Fig S1B. (C) Sagittal view of alcian blue/ alizarin red stained mouse skull at E16.5 for different genotypes. Frontal (F) and nasal (N) bone, mandible (MD), premaxilla (PMX), maxilla (MX) and the squamous (SQ) bone was severely affected in *Fgfr1*^*cKO/cKO*^; *Fgfr2*^*cKO/cKO*^ conditional double mutants. In *Fgfr1*^*cKO/cKO*^; *Fgfr2*^*cKO*/+^ mutants also showed partial mandibular defect where proximal condylar (CDP), coronoid (CRP) and angular process (AGP) of the mandible was affected. (D) Alcian blue/ alizarin red staining of mouse skulls at E14.5. Alizarin red staining mark ossified region in mandible and maxilla in control (*Fgfr1^cKO/+^; Fgfr2^cKO/+^)* skull. Yellow arrowhead marks the Meckel’s cartilage. In *Fgfr1*^*cKO/cKO*^; *Fgfr2*^*cKO*/+^ mutants, we observed reduced ossification (red arrow) in the mandible with a significantly smaller Meckel’s cartilage (yellow arrow). Meckel’s cartilage was lost in *Fgfr1/2* conditional double mutant with no observable ossification.

**Extended Data Fig. 2:**
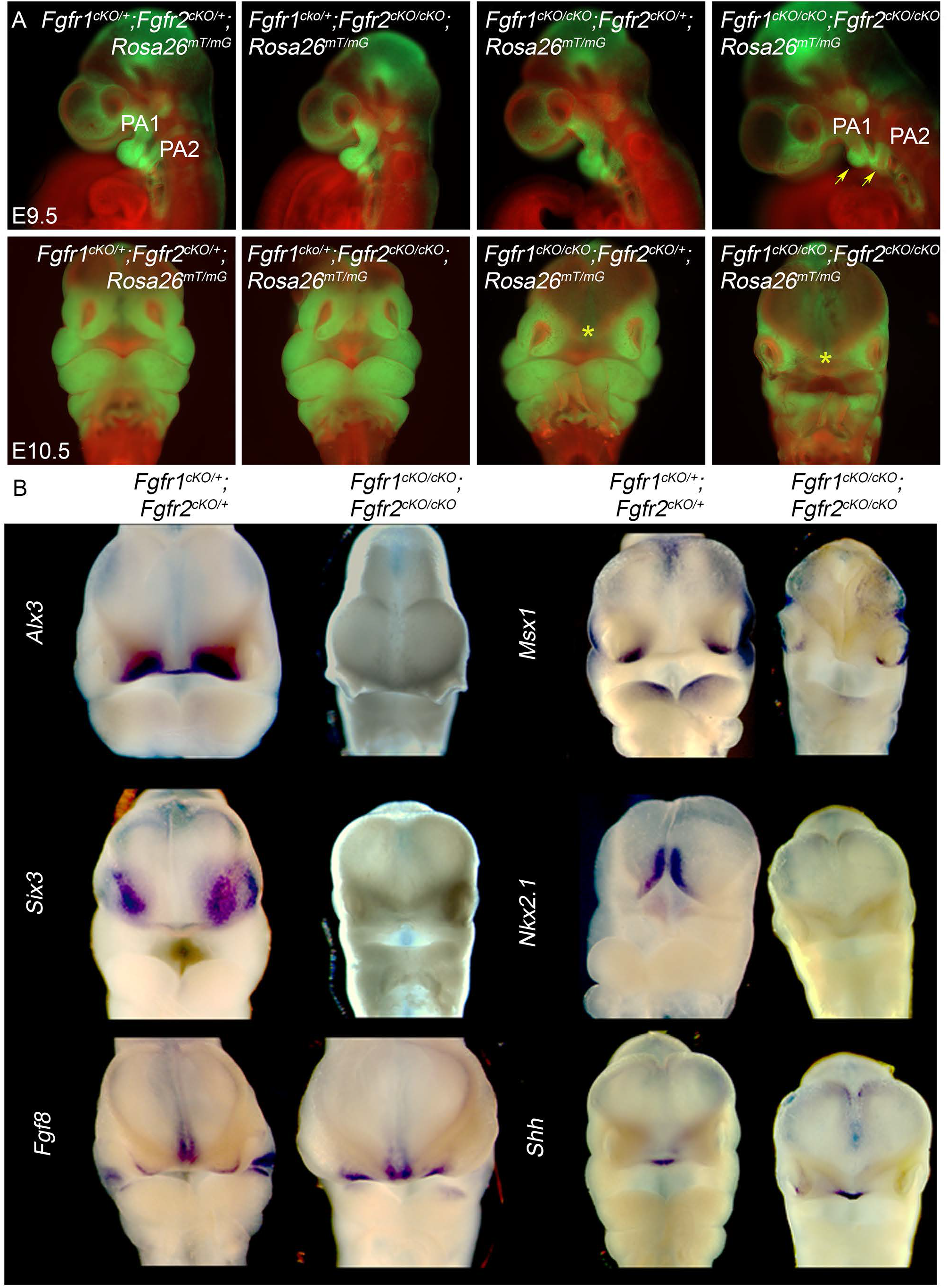
Molecular changes and defects in craniofacial ossification in *Fgfr1*^*cKO/cKO*^; *Fgfr2*^*cKO/cKO*^ double mutants. (A) Conditional *Fgfr1*-*Fgfr2* null mutant embryos carrying the *ROSA26*^*mT/mG*^ reporter analyzed in whole mount. cNCCs in the head express GFP (green). At E9.5 (top panel), *Fgfr1/2* conditional double mutants (*Fgfr1*^*cKO/cKO*^; *Fgfr2*^*cKO/cKO*^; *ROSA26*^*mT/mG*^) had hypoplastic pharyngeal arches PA1 and PA2 (yellow arrows), shown by reduced GFP fluorescence compared to *Fgfr1*-*Fgfr2* conditional heterozygotes. *Fgfr1^cKO/+^; Fgfr2^cKO/cKO^* and *Fgfr1*^*cKO/cKO*^; *Fgfr2*^*cKO*/+^ mutants appeared normal at this stage. Frontal view at E10.5 (lower panels). A wide midline separation was observed in both *Fgfr1/2* conditional double (*Fgfr1*^*cKO/cKO*^; *Fgfr2*^*cKO/cKO*^; *ROSA26*^*mT/mG*^) and *Fgfr1*^*cKO/cKO*^; *Fgfr2*^*cKO*/+^; *ROSA26*^*mT/mG*^ mutants (yellow asterisk) compared to controls and *Fgfr1^cKO/+^; Fgfr2^cKO/cKO^*; *ROSA26*^*mT/mG*^ mutants. (B) Whole mount mRNA *in situ* hybridization was used to compare the expression of *Alx3, Msx1, Six3, Nkx2.1, Fgf8* and *Shh* at E10.5 in the facial primordia for *Fgfr1^cKO/+^; Fgfr2^cKO/+^* control and *Fgfr1*^*cKO/cKO*^; *Fgfr2*^*cKO/cKO*^ mutants. *Msx1* expression was reduced, and *Alx3*, *Six3* and *Nkx2.1* expression were lost in *Fgfr1/2* conditional double mutants. The expression of *Fgf8* and *Shh* remained unaffected in the *Fgfr1*^*cKO/cKO*^; *Fgfr2*^*cKO/cKO*^ mutants.

**Extended Data Fig. 3:**
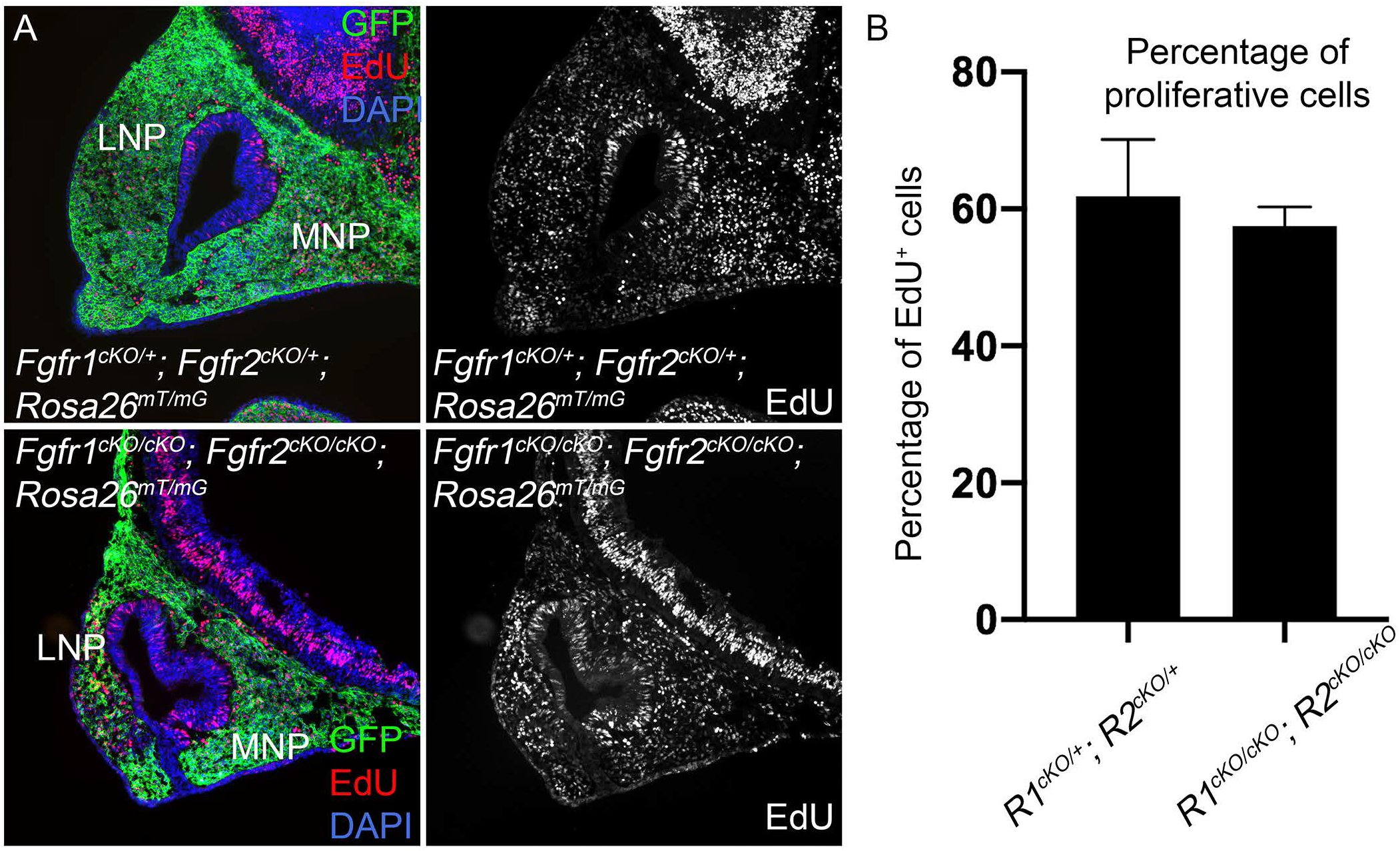
Cell proliferation in *Fgfr1*^*cKO/cKO*^; *Fgfr2*^*cKO/cKO*^ double mutants. (A) EdU incorporation was used to assay cell proliferation in *Fgfr1^cKO/cKO^; Fgfr2^cKO/cKO^; ROSA26^mT/mG^* double mutants. No change in the proliferation index of GFP^+^ cNCC lineage cells was observed at E10.5 in the MNP and LNP across various genotypes. (B) Quantification of the mean percentage of EdU^+^ cells normalized to GFP^+^ cNCC lineage cells, in control versus double null mutant cells suggests cell proliferation remained unaffected in *Fgfr1*^*cKO/cKO*^; *Fgfr2*^*cKO/cKO*^ double mutants.

**Extended Data Fig. 4:**
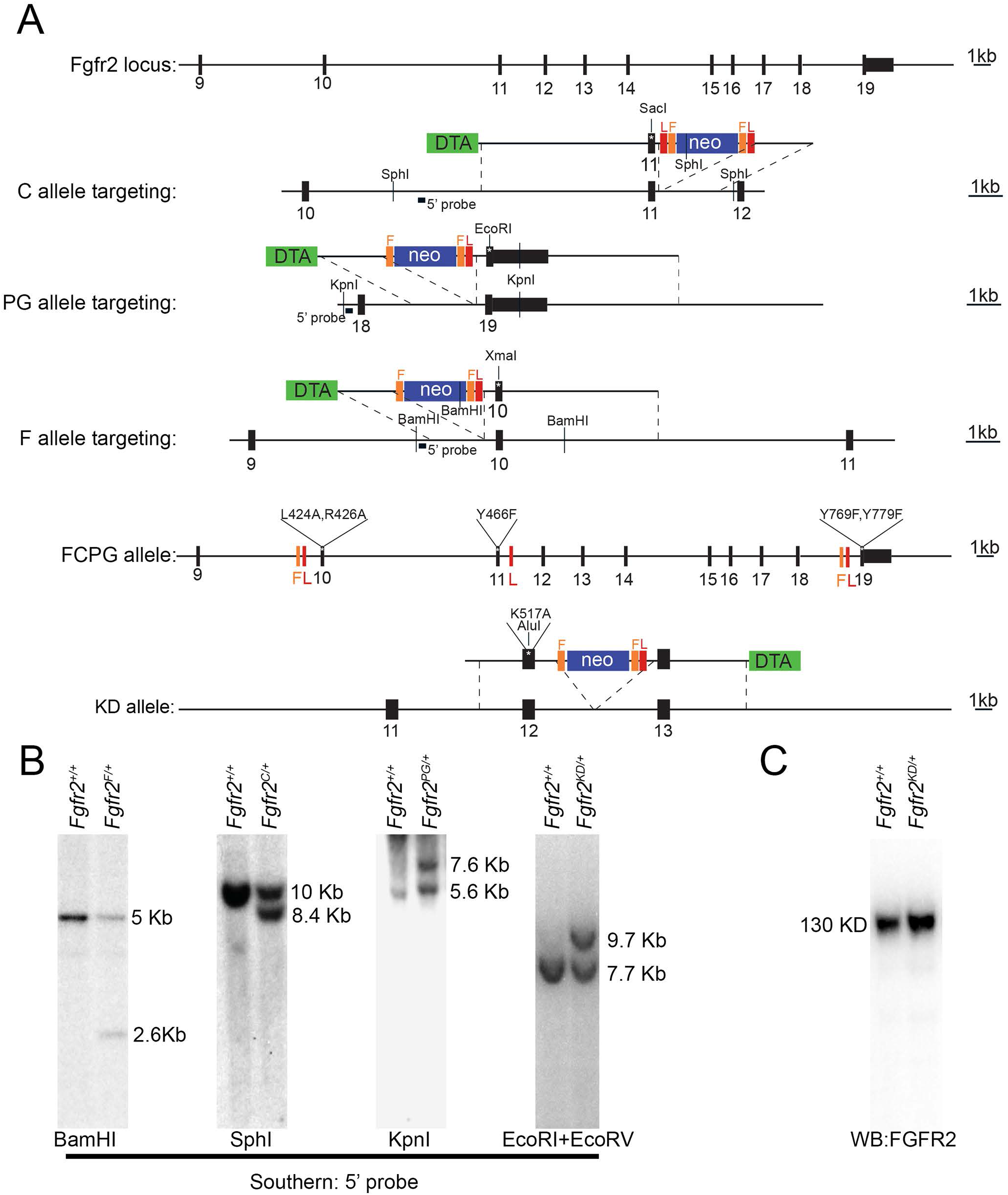
Allelic series of *Fgfr2* signaling mutations. (A) Representation of *Fgfr2* WT locus, targeting vectors and the corresponding targeted loci for C, PG and F alleles, the *Fgfr2*^*FCPG*^ locus, and the KD allele. Targeting vectors C, PG and F introduced Y466F, Y769F/Y779F and L434A/R426A mutations sequentially to the *Fgfr2* locus, respectively. The targeting vector for KD allele introduced the K517A mutation. ES cells were targeted first with C targeting vector generating *Fgfr2^+/C^* mutant cells. *Fgfr2^+/C^* ES cells were targeted with PG targeting vector generating either *Fgfr2^C/PG^* or *Fgfr2^+/CPG^* mutant cells. Finally, *Fgfr2^+/CPG^* was targeted with F targeting vector resulting in *Fgfr2^F/CPG^* or *Fgfr2^+/FCPG^* mutant ES cells. Multiple LoxP sites were introduced at the *Fgfr2* locus during successive rounds of targeting, and the neo cassette was removed *in vivo* or in culture before generating signaling mouse mutants. We were unable to perform a conditional mutation for the *Fgfr2*^*FCPG*^ allele due to the serial retention of multiple lox sites during the generation of this allele. Abbreviation: L, loxP site; F, FRT site; DTA, Diptheria toxin A cassette; Neo, Neomycin resistance cassette. (B) Southern blot confirmation for proper targeting of F, C, PG and KD mutations at the *Fgfr2* locus. 5’ probes used for each targeting events are indicated. BamHI restriction enzyme generated 5kb and 2.6kb band for WT and F targeted loci, respectively. SphI digestion generated 10kb and 8.4 kb fragment for WT and C targeted loci, respectively. KpnI digestion generated a 5.6kb band for WT locus and a 7.6kb band for PG targeted locus. EcoRI and EcoRV restriction digestion generated 7.7kb WT fragment and 9.7kb KD-mutant fragment. (C) Western blot from facial prominence cells harvested from E9.5 wild-type and *Fgfr2^KD/KD^* embryos show full length FGFR2^KD^ protein expression.

**Extended Data Fig. 5:**
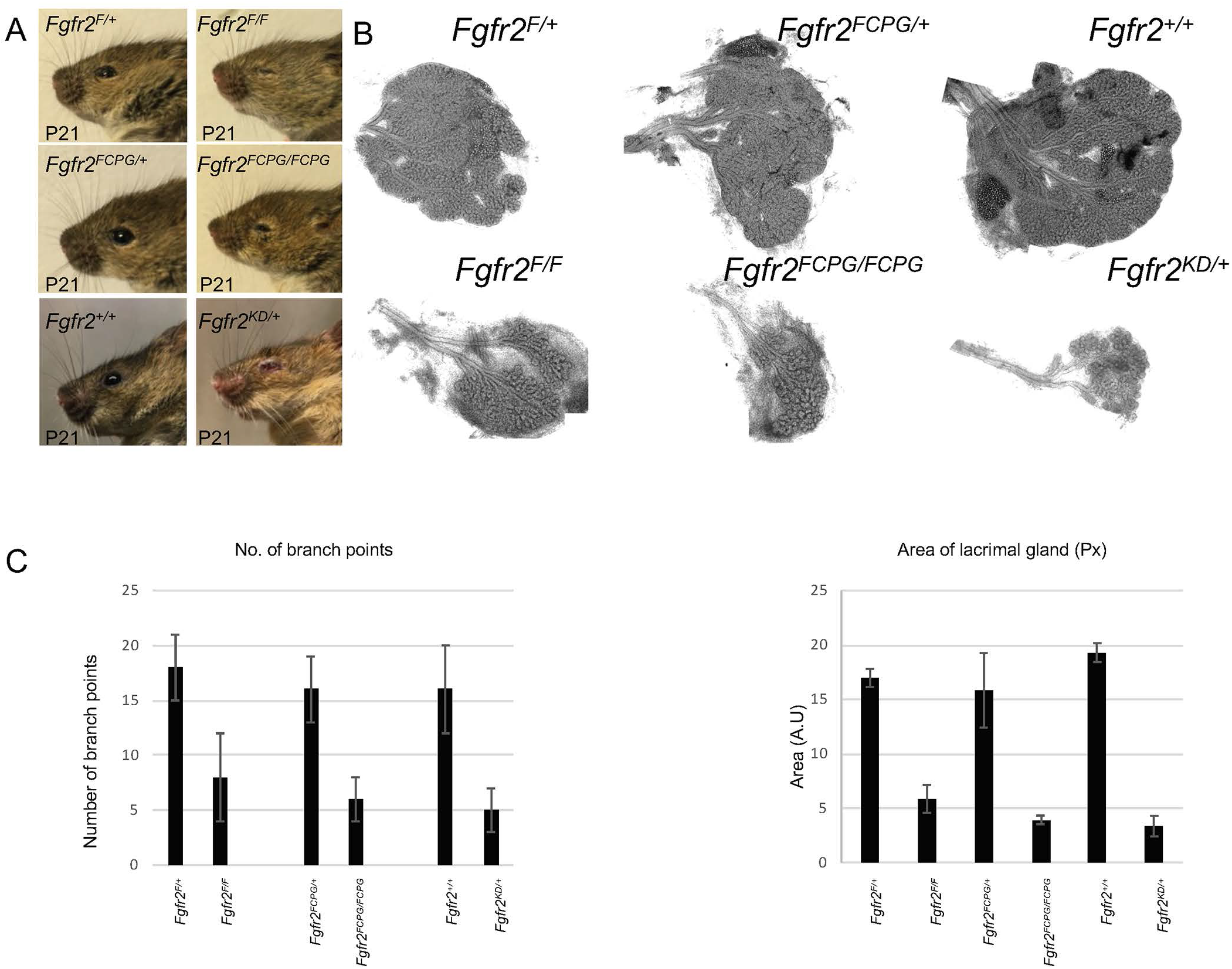
Lacrimal gland defects in *Fgfr2* signaling mutants. (A) Sagittal view of the left eye at P21 for the mentioned genotypes. *Fgfr2*^*F*/*F*^, *Fgfr2*^*FCPG/FCPG*^ and *Fgfr2*^*KD*/+^ mice showed fully penetrant defects with onset at P15 which included laceration around the eye. More severe defects were observed in *Fgfr2*^*KD*/+^ mutants. (B) The lacrimal gland from signaling mutants was dissected at P5 and stained with acetocarmine. Representative images are shown here for *Fgfr2*^*F/F*^, *Fgfr2*^*FCPG/FCPG*^ and *Fgfr2*^*KD*/+^ compared to littermate *Fgfr2*^*F*/+^, *Fgfr2*^*FCPG*/+^ and *Fgfr2*^+/+^ controls. (C) Acetocarmine stained preparation of exorbital lacrimal glands at P5 from *Fgfr2*^*F/F*^, *Fgfr2*^*FCPG/FCPG*^ and *Fgfr2*^*KD*/+^ mice showed reduced branching and reduced overall size of the gland. Flat mounts of the acetocarmine stained lacrimal glands for each genotype were microscopically analyzed and primary and secondary branching points were counted manually. The bar graph shows the mean ± standard deviation of the relative number of branch points for each genotype analyzed. The relative size of the lacrimal gland for each genotype was also assessed by comparing images acquired in the same magnification as a mean of the area observed in flat mounts. *Fgfr2*^*F/F*^, *Fgfr2*^*FCPG/FCPG*^ and *Fgfr2*^*KD*/+^ compared to littermate *Fgfr2*^*F*/+^, *Fgfr2*^*FCPG*/+^ and *Fgfr2*^+/+^ controls show reduced branching and a reduction in the overall size of the gland.

**Extended Data Fig. 6:**
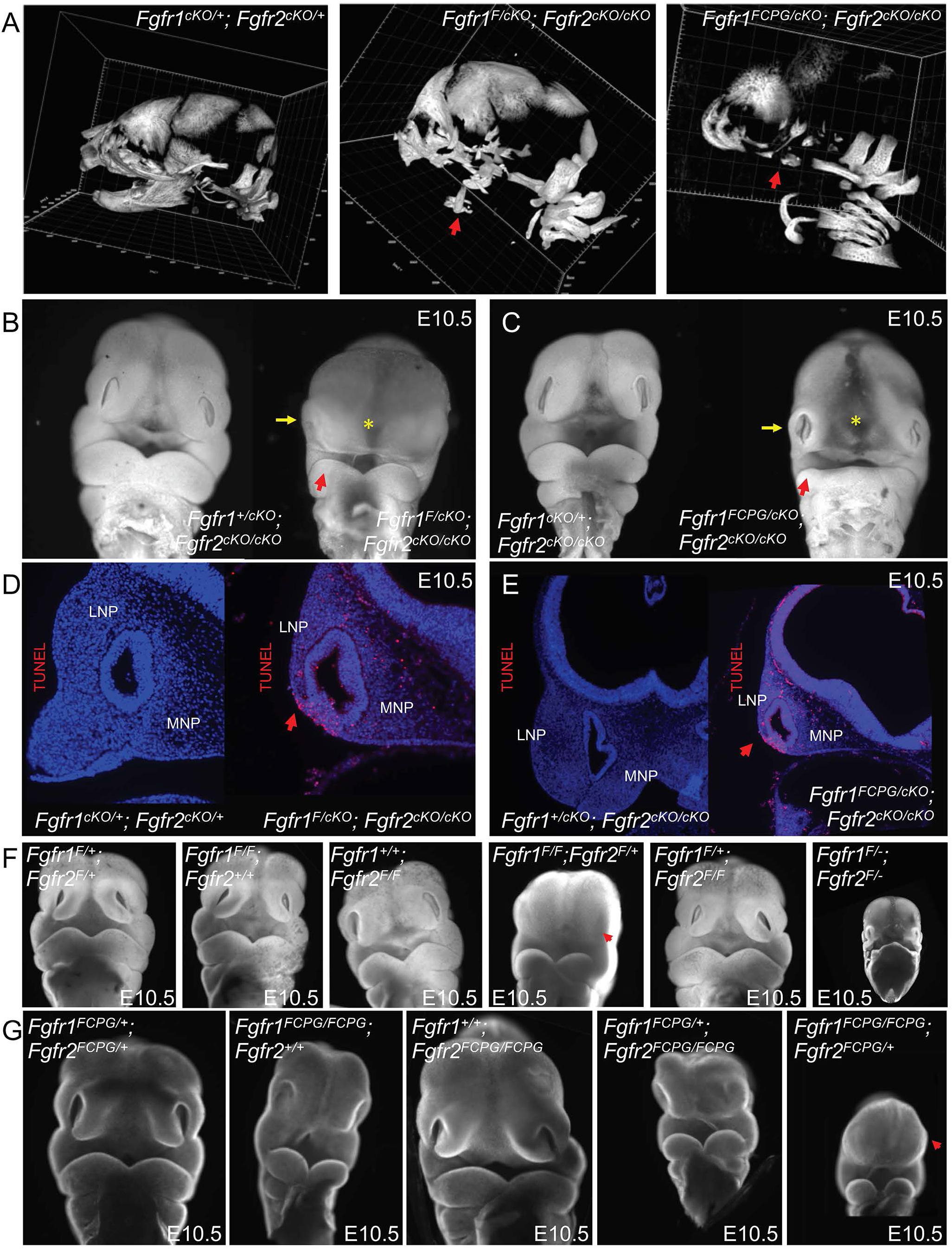
Analysis of *Fgfr1* and *Fgfr2* single and compound signaling mutants. (A) Micro-CT analysis of control *Fgfr1*^*cKO*/+^; *Fgfr2*^*cKO*/+^ embryos at E18.5 compared to *Fgfr1*^*F/cKO*^; *Fgfr2*^*cKO1*/+^ and *Fgfr1*^*FCPG/cKO*^; *Fgfr2*^*cKO*/+^ embryos at similar stages. The mandible (red arrow) was more severely affected in *Fgfr1*^*FCPG/cKO*^; *Fgfr2*^*cKO*/+^ than in *Fgfr1*^*F/cKO*^; *Fgfr2*^*cKO*/+^ embryos. (B) Frontal facial view of DAPI-stained E10.5 *Fgfr1*^*F/cKO*^; *Fgfr2*^*cKO*/+^ embryos compared to control *Fgfr1*^*cKO*/+^; *Fgfr2*^*cKO*/+^ embryos. Midline separation (yellow asterisk) was observed and facial prominences were hypoplastic (yellow arrow), however, no defect in the mandible (red arrow) was observed. (C) Frontal facial view of DAPI-stained E10.5 *Fgfr1*^*FCPG/cKO*^; *Fgfr2*^*cKO*/+^ embryos compared to control *Fgfr1*^*cKO*/+^; *Fgfr2*^*cKO*/+^ embryos. In the mutants, midline separation (yellow asterisk) and facial prominence (yellow arrow) defects were more severe than *Fgfr1*^*F/cKO*^; *Fgfr2*^*cKO*/+^ embryos. A defect in the mandible (red arrow) was also more severe than *Fgfr1*^*F/cKO*^; *Fgfr2*^*cKO*/+^ embryos. (D-E) Cell-death analysis by TUNEL staining in the facial prominences of (D) *Fgfr1*^*F/cKO*^; *Fgfr2*^*cKO*/+^ and (E) *Fgfr1*^*FCPG/cKO*^; *Fgfr2*^*cKO*/+^ embryos at E10.5, showing multiple dying cells in both the MNP and the LNP compared to the *Fgfr1*^*cKO*/+^; *Fgfr2*^*cKO*/+^ controls. (F) Frontal facial view of DAPI-stained *Fgfr1*^*F*^ and *Fgfr2*^*F*^ compound mutant embryos at E10.5. *Fgfr1*^*F/F*^; *Fgfr2*^+/*F*^ embryos had defects in the development of facial prominences, as opposed to *Fgfr1^F/+^; Fgfr2^F/+^* controls. Mandibular prominences were hypoplastic, however, the maxillary prominences, FNP and MNP were more severely affected. *Fgfr1*^*F/F*^; *Fgfr2*^+/+^ mutants, *Fgfr1*^+/+^; *Fgfr2*^*F/F*^ mutants, and *Fgfr1*^*F*/+^; *Fgfr2*^*F/F*^ mutants did not show obvious defects at this stage. *Fgfr1*^*F/F*^; *Fgfr2*^+/*F*^ mutant embryos were also developmentally retarded. We did not recover *Fgfr1*^*F/F*^; *Fgfr2*^*F/F*^ mutant embryos by E9.5. (G) The craniofacial phenotype at E10.5 of DAPI-stained *Fgfr1*^*FCPG*^ and *Fgfr2*^*FCPG*^ compound mutant embryos. We observed the posterior truncation for *Fgfr1^FCPG/FCPG^* mutant embryos. The introduction of one copy of *Fgfr2*^*FCPG*^ allele on a *Fgfr1^FCPG/FCPG^* background resulted in severe growth retardation of the facial prominences compared to control *Fgfr1^+/FCPG^; Fgfr2^FCPG/FCPG^* embryos. *Fgfr2^FCPG/FCPG^* and *Fgfr1^+/FCPG^; Fgfr2^FCPG/FCPG^* embryos did not exhibit abnormal phenotypes and were comparable to WT embryos.

**Extended Data Fig. 7:**
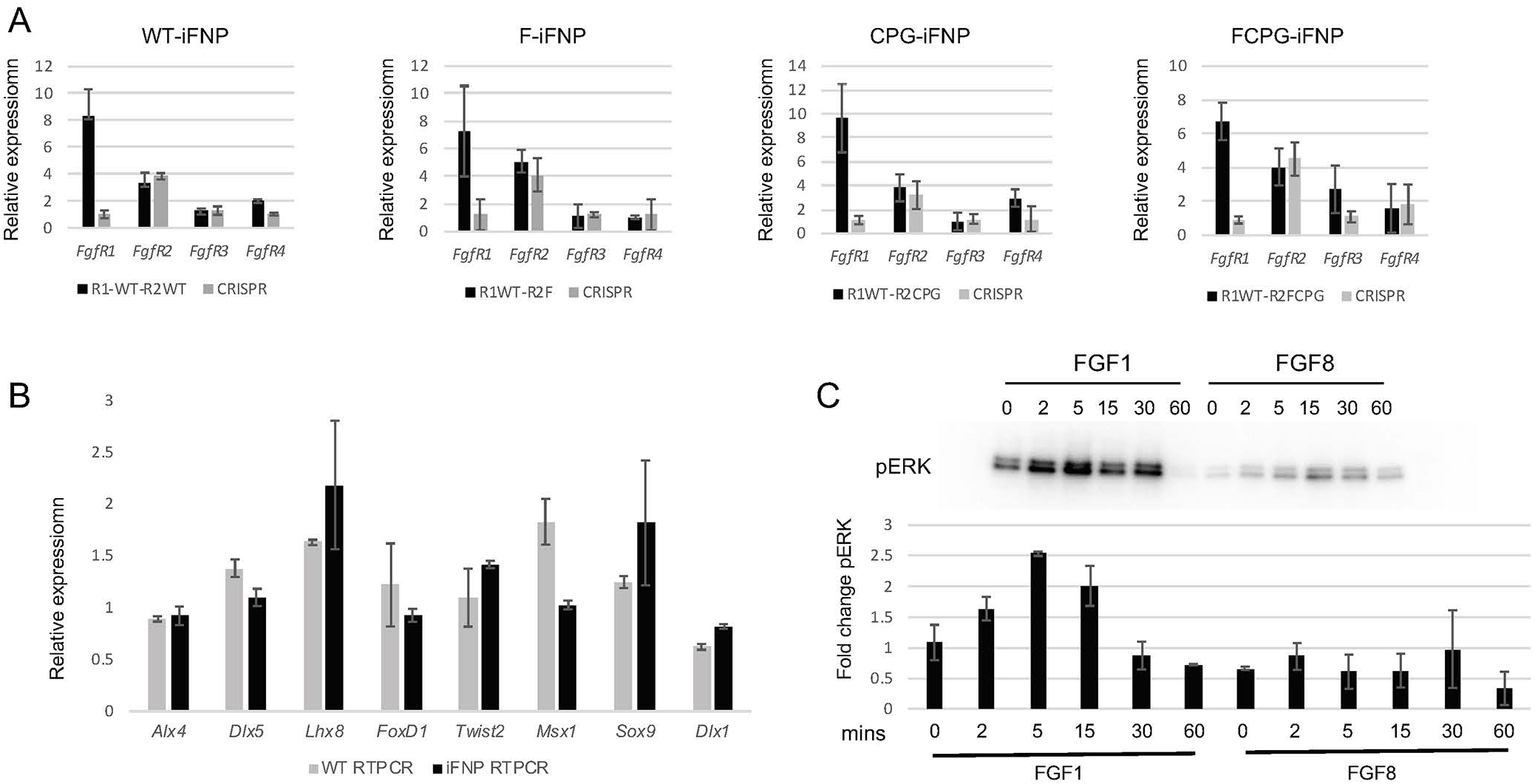
Characterization of *Fgfr1* null mutant iFNP cells. (A) *Fgfr1, Fgfr3, Fgfr4* and *Fgfr2*^*WT*^ mRNA expression in wild-type and *Fgfr2*^*F*^, *Fgfr2*^*CPG*^ and *Fgfr2*^*FCPG*^ iFNP cells before and after *Fgfr1*-CRISPR disruption, as determined by RT-qPCR. (B) *Alx4, Dlx1, Dlx5, FoxD1, Twist1* and *Lhx8* were all expressed at similar levels by quantitative real-time PCR (qRT-PCR) in both immortalized FNP (iFNP) cells and primary FNP cells derived from wild-type embryos. Only *Msx1* expression was 1.8-fold higher in wild-type primary FNP cells compared to iFNP cell lines. These results identify iFNP cells as a suitable cell type to interrogate FGF signaling functions in craniofacial development. (C) *WT*-iFNP cells were serum starved overnight and stimulated with either 50 ng/mL FGF1 or 50 ng/mL FGF8-b and 5 μg/mL heparin for the indicated times (0-60 mins). Robust ERK1/2 activation was observed with FGF1 treatment compared to FGF8-b treatment.

**Extended Data Fig. 8:**
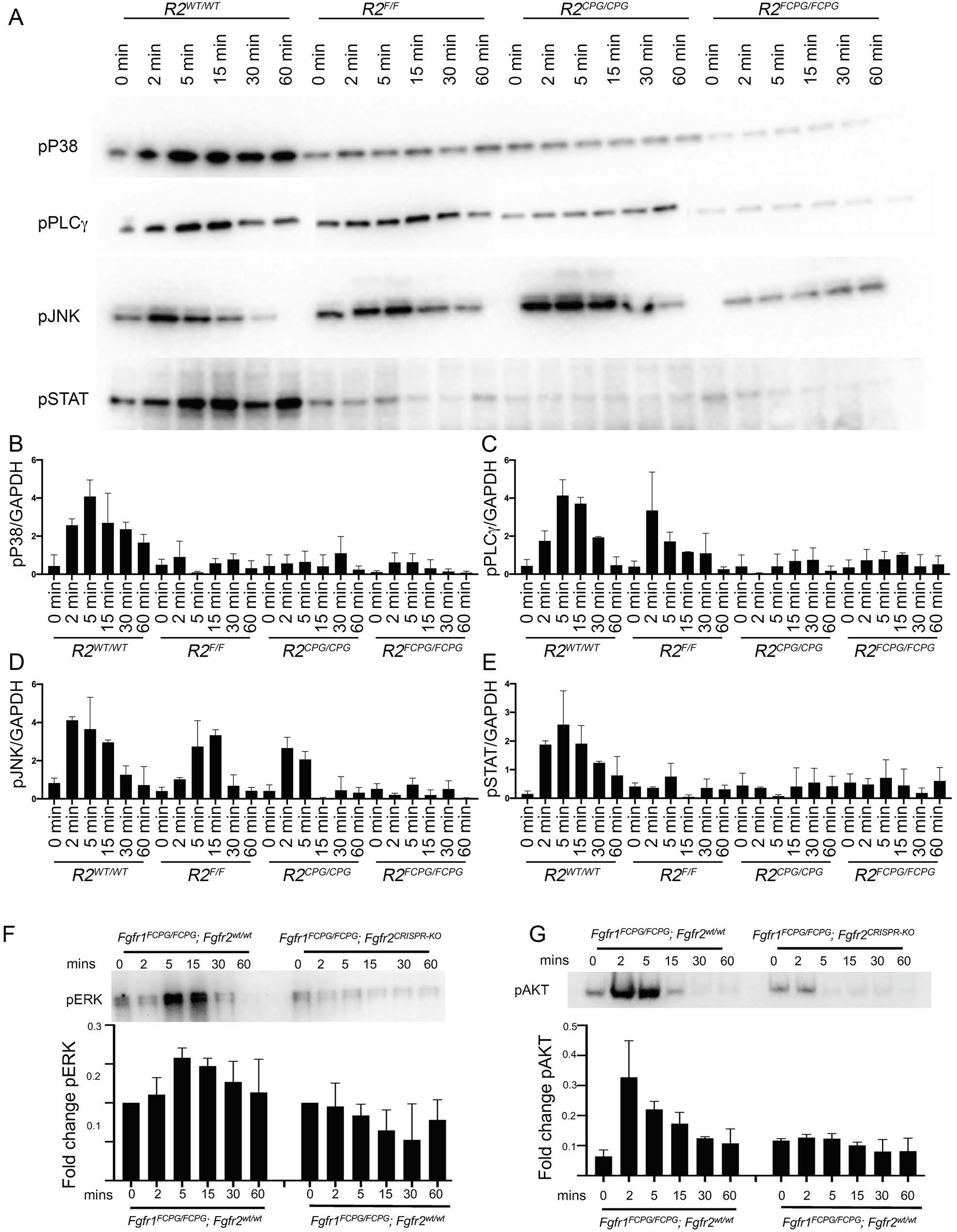
Intracellular signaling upon FGF1 stimulation. (A) iFNP cells derived from the indicated genotypes, *Fgfr2*^*wt/wt*^ (*R2*^*wt/w*^), *Fgfr2*^*F/F*^ (*R2*^*F/F*^), *Fgfr2*^*CPG/CPG*^ (*R2*^*CPG/CPG*^) and *Fgfr2*^*FCPG/FCPG*^ (*R2*^*FCPG/FCPG*^), were serum starved overnight and stimulated with 50 ng/mL FGF1 and 5 μg/mL heparin for the indicated times (0-60 mins). Phospho-blots were stripped and reblotted with GAPDH as a loading control. Activation of pP38, pPLCγ, pJNK and pSTAT3 was investigated. (B-E) Quantification of pathway activation for (B) pP38, (C) pPLCγ, (D) pJNK and (E) pSTAT3 normalized to the GAPDH loading control, is reported as mean ± standard deviation with a minimum of three independent biological replicates. (F-G) *Fgfr1*^*FCPG*^ *Fgfr2* null iFNP (*Fgfr1*^*FCPG*^; *Fgfr2*^*CRISPR-KO*^) cells generated using CRISPR/Cas9. Upon FGF1 stimulation, we observed peak activation of (F) pERK1/2 after 2 mins in *Fgfr1*^*FCPG/FCPG*^ iFNP cells. We did not observe activation over background in *Fgfr1*^*FCPG/FCPG*^; *Fgfr2*^*CRISPR-KO*^ iFNP cells. (G) pAKT activation was undetectable in both *Fgfr1*^*FCPG/FCPG*^; *Fgfr2*^*CRISPR-KO*^ in contrastto *Fgfr1*^*FCPG/FCPG*^; *Fgfr2*^+/+^ iFNP cells.

**Extended Data Fig. 9:**
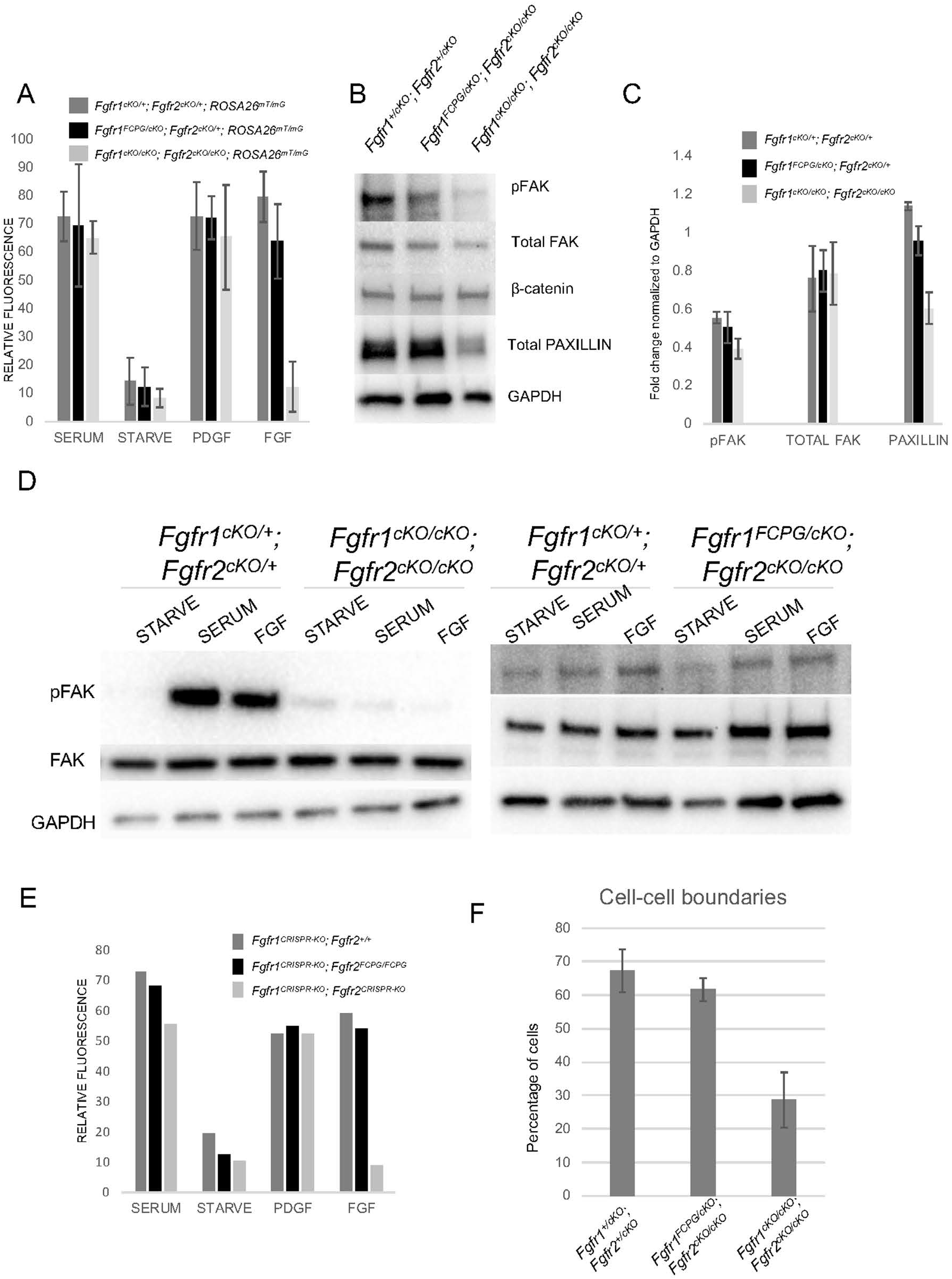
FGFR signaling in cell-matrix and cell-cell adhesion. (A) Quantification of Paxillin immunostaining mean fluorescence intensity in FNP cells, shown for different genotypes in Fig. 4B. Mean fluorescence intensity in starvation media act as a baseline. Upon serum stimulation or PDGF treatment, cells from all genotypes showed a robust increase in focal adhesions. Upon FGF treatment, control cells and *Fgfr1*^*FCPG/cKO*^; *Fgfr2*^*cKO*/+^; *ROSA26*^*mT/mG*^ FNP cells showed robust paxillin accumulation, in contrast to *Fgfr1*^*cKO/cKO*^; *Fgfr2*^*cKO/cKO*^; *ROSA26*^*mT/mG*^ double mutant cells. (B) Western Blot analysis of freshly harvested FNP cells from control, *Fgfr1*^*FCPG/cKO*^; *Fgfr2*^*cKO*/+^; *ROSA26*^*mT/mG*^ and *Fgfr1*^*cKO/cKO*^; *Fgfr2*^*cKO/cKO*^; *ROSA26*^*mT/mG*^ double mutant FNP cells for pFAK, total FAK, β-catenin and Paxillin. GAPDH was used as a loading control. Total FAK and β-catenin levels remained constant across all genotypes. (C) Bar graph shows relative levels of pFAK, total FAK and Paxillin. pFAK and Paxillin levels were comparable in control and *Fgfr1*^*FCPG/cKO*^; *Fgfr2*^*cKO*/+^; *ROSA26*^*mT/mG*^ FNP cells, but severely reduced in *Fgfr1*^*cKO/cKO*^; *Fgfr2*^*cKO/cKO*^; *ROSA26*^*mT/mG*^ FNPs. Total FAK levels remained unchanged. (D) Freshly harvested *Fgfr1*^*cKO*/+^; *Fgfr2*^*cKO*/+^; *ROSA26*^*mT/mG*^ control, *Fgfr1*^*FCPG/cKO*^; *Fgfr2*^*cKO*/+^; *ROSA26*^*mT/mG*^, and *Fgfr1*^*cKO/cKO*^; *Fgfr2*^*cKO/cKO*^; *ROSA26*^*mT/mG*^ double mutant FNP cells were treated with serum or FGF1 for 15 mins, before assessing levels of pFAK by western blot. *Fgfr1*^*cKO*/+^; *Fgfr2*^*cKO*/+^; *ROSA26*^*mT/mG*^ control and *Fgfr1*^*FCPG/cKO*^; *Fgfr2*^*cKO*/+^; *ROSA26*^*mT/mG*^ FNP cells showed a comparable increase in FAK activation upon FGF stimulation. In contrast, *Fgfr1*^*cKO/cKO*^; *Fgfr2*^*cKO/cKO*^; *ROSA26*^*mT/mG*^ double mutant FNP cells failed to activate FAK in both serum-stimulated and FGF treated conditions. (E) Quantification of mean fluorescence from Paxillin immunostaining in *Fgfr2*^*WT/WT*^ (*Fgfr1CRISPR-KO Fgfr2WT/WT*), *Fgfr2FCPG/FCPG* (*Fgfr1CRISPR-KO Fgfr2FCPG/FCPG*), and *Fgfr2*^−/−^ (*Fgfr1CRISPR-KO Fgfr2*^*CRISPR-KO*^)-iFNP cells upon serum starvation and growth factor treatments shown in Fig. 4B. Upon serum stimulation or PDGF treatment, cells from all genotypes showed a robust increase in focal adhesions. Upon FGF treatment, both control and *Fgfr1*^*CRISPR-KO*^ *Fgfr2*^*FCPG/FCPG*^ iFNP cells showed robust paxillin accumulation at focal adhesions, whereas *Fgfr1*^*CRISPR-KO*^ *Fgfr2 CRISPR-KO* double null mutant iFNP cells showed poor response. (F) Quantification of percentage of cells that form cell boundaries (n=500 contacts for each genotype) in *Fgfr1*^*cKO*/+^; *Fgfr2*^*cKO*/+^; *ROSA26*^*mT/mG*^ control, *Fgfr1*^*FCPG/cKO*^; *Fgfr2*^*cKO*/+^; *ROSA26*^*mT/mG*^, and *Fgfr1*^*cKO/cKO*^; *Fgfr2*^*cKO/cKO*^; *ROSA26*^*mT/mG*^ double mutant FNP cells. When freshly harvested primary FNP cells were serum stimulated for 3 hrs, both control and *Fgfr1*^*FCPG/cKO*^; *Fgfr2*^*cKO*/+^; *ROSA26*^*mT/mG*^ cells formed comparable numbers of cell-cell contacts (Figure 7D). In contrast, this number was reduced to half in double mutant FNP cells.

**Supplementary Information Figure S1:**
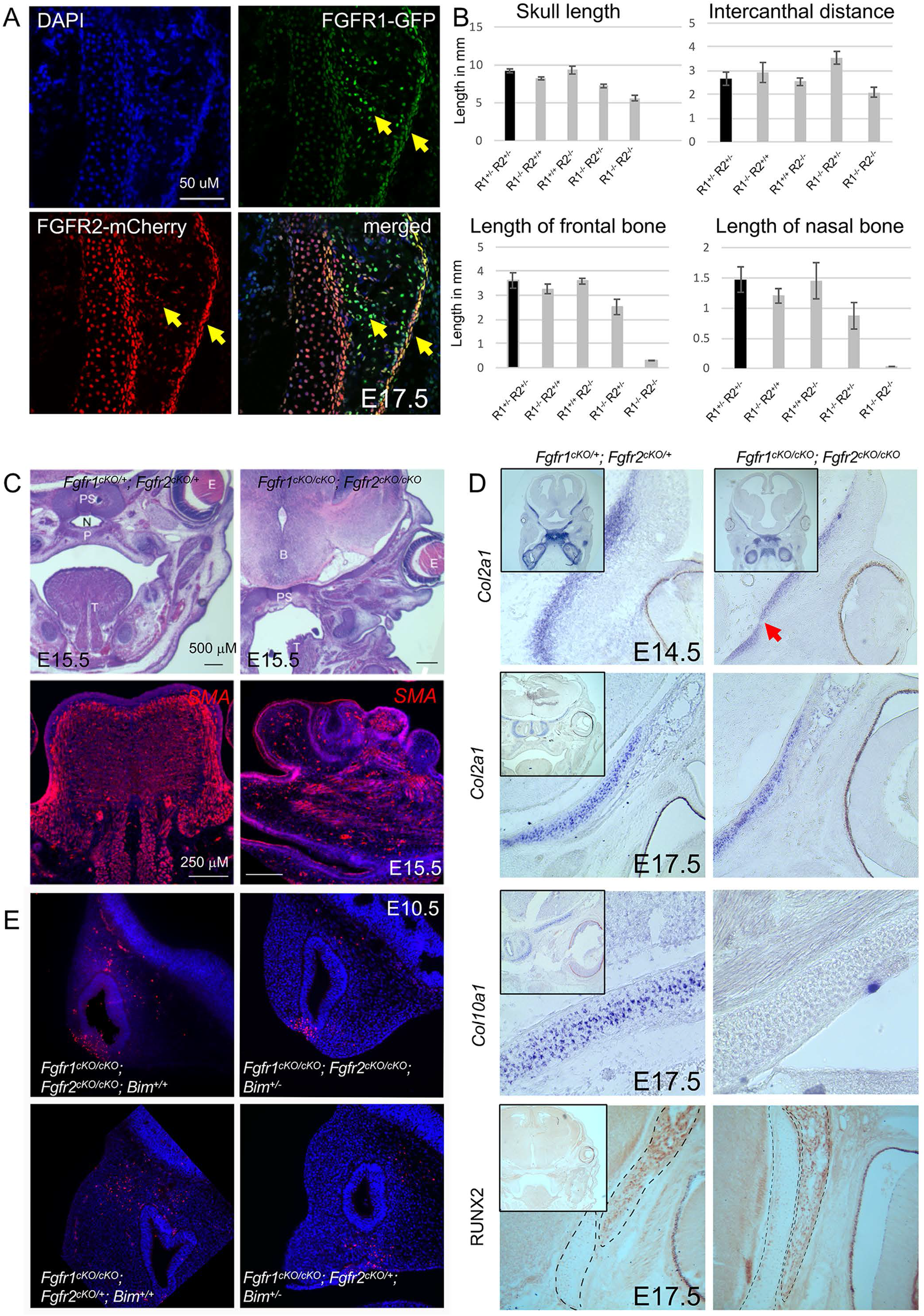
Craniofacial ossification and organogenesis defects in *Fgfr1/2* mutants. (A) Spatial domain of FGFR1 and FGFR2 expression in the frontal bone at E17.5. GFP and mCherry immunohistochemistry was used to detect expression from *Fgfr1-GFP* and *Fgfr2-mCherry* reporter alleles. Co-expression of both FGFR1-GFP and FGFR2-mCherry was observed in the cartilage and the perichondrium (yellow arrow). (B) Quantification of skull length, intercanthal distance, length of frontal and nasal bone for different genotypes measured in millimeters (mm). (C) Histological examination at E15.5 showed defective organogenesis in multiple organs, including the palate, tongue, and skeleton. In the double mutants several skeletal structures such as palate (P), nasopharyngeal lumen (N) and pre-sphenoid (PS) were affected compared to the controls. Non-neural crest derived muscles in the tongue were also affected. SMA immunofluorescence showed organized muscle fiber differentiation in controls. In the *Fgfr1/2* mutants, muscle fiber differentiation appeared random and disorganized. (D) Differentiation of craniofacial skeleton was assessed by studying the expression of cartilage differentiation marker, *Col2a1*, on sections. mRNA *in situ* hybridization analysis at E14.5 showed *Col2a1* expression in the chondrocranium (skull base) and mandible in the control (inset shows low magnification view of the region). In the double mutants, however, expression was restricted to a narrow band of cells in the skull base. Expression of *Col2a1* and of *Col10a1* and Runx2 (mature cartilage markers) was assessed at E17.5 on sections (inset shows a low magnification view of the region). *Col2a1* mRNA expression was restricted to the skull base, mandible and frontal bone in controls. A similar expression was also observed in conditional double mutants. At E17.5, *Col10a1* mRNA expression was much broader in the controls. In conditional double mutants, *Col10a1* expression was undetectable suggesting a block in terminal chondrogenic differentiation. We detected the expression of mature cartilage and late osteoblast marker, RUNX2, in both controls and the conditional double mutants. **(E)** Increased cell-death observed in conditional double mutants was partially rescued in *Fgfr1*^*cKO/cKO*^; *Fgfr2*^*cKO/cKO*^; *Bim*^+/−^ mutants at E10.5. More TUNEL^+^ cells were observed in the LNP in *Fgfr1*^*cKO/cKO*^; *Fgfr2*^*cKO/cKO*^ and *Fgfr1*^*cKO/cKO*^; *Fgfr2*^*cKO*/+^ mutants compared to corresponding *Fgfr1*^*cKO/cKO*^; *Fgfr2*^*cKO/cKO*^; *Bim*^+/−^ and *Fgfr1*^*cKO/cKO*^; *Fgfr2*^*cKO*/+^; *Bim*^+/−^ counterparts. Compared to controls, reduced TUNEL positive foci were observed in *Fgfr1*^*cKO/cKO*^; *Fgfr2*^*cKO/cKO*^; *Bim*^+/−^ mutants.

**Supplementary Information Figure S2:**
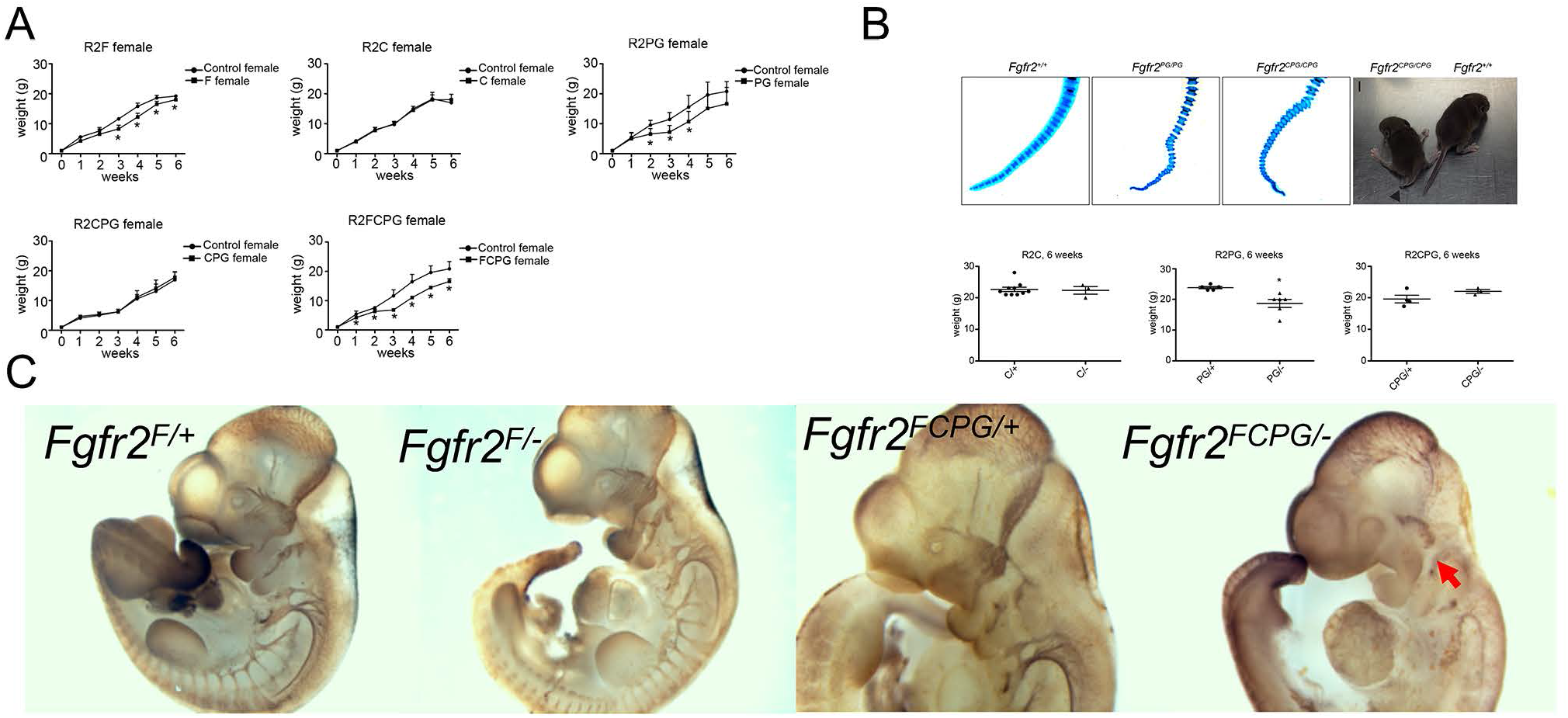
Growth defects in *Fgfr* signaling mutants. (A) Growth rate (in terms of weight gain) of homozygous *Fgfr2*^*F/F*^, *Fgfr2*^*C/C*^, *Fgfr2*^*PG/PG*^, *Fgfr2*^*CPG/CPG*^, and *Fgfr2*^*FCPG/FCPG*^ female versus control until 6 weeks shown as mean ± SEM. * indicates p-value <0.05. We noted a decrease in growth rate for *Fgfr2*^*F/F*^, *Fgfr2*^*PG/PG*^, and *Fgfr2*^*FCPG/FCPG*^ mutant mice compared to control. (B) *Fgfr2*^*PG/PG*^ and *Fgfr2*^*CPG/CPG*^ skeletal preparation at P0 showed fusion of the caudal vertebrae, resulting in a kinked or curly tail phenotype in rare cases (black arrow). Hemizygote *Fgfr2*^*C*/−^, *Fgfr2*^*PG*/−^ and *Fgfr2*^*CPG*/−^ mice were weighed at 6 weeks. *Fgfr2*^*PG*/−^ were smaller than their control littermate. (C) Whole mount immunohistochemistry for neurofilament staining revealed trigeminal nerve defects at E10.5 in *Fgfr2*^*FCPG*/−^ embryo but not in *Fgfr2*^*F*/−^. In *Fgfr2*^*F*/+^, *Fgfr2*^*FCPG*/+^ and *Fgfr2*^*F*/−^ embryos, trigeminal nerve migrated to the anterior domain of the PA1 (red arrow) at E10.5. Trigeminal nerves fail to migrate into the pharyngeal arch of *Fgfr2*^*FCPG*/−^ mutants.

**Supplementary Information Table 1:** Recovery of *Fgfr2* homozygous and *Fgfr2* hemizygous signaling mutants. *Fgfr2* signaling mutant heterozygotes (*Fgfr2*^*F*/+^, *Fgfr2*^*C*/+^, *Fgfr2*^*PG*/+^, *Fgfr2*^*CPG*/+^, *Fgfr2*^*FCPG*/+^ and *Fgfr2*^*KD*/+^) were intercrossed. Embryos were analyzed either at E18.5 or postnatally between P4 and P10. To identify *in vivo* phenotypes associated with *Fgfr2* signaling mutations, *Fgfr2* signaling mutant heterozygotes were crossed with *Fgfr2*^+/−^ null heterozygotes. *Fgfr2*^*C*/−^, *Fgfr2*^*PG*/−^ and *Fgfr2*^*CPG*/−^ embryos were analyzed at E18.5 and postnatally at P0 and between P4 and P10. Hemizygous *Fgfr2*^*F*/−^ and *Fgfr2*^*FCPG*/−^ mutant mice were recovered at expected Mendelian ratios at E18.5, but died postnatally.

| Recovery of *Fgfr2* signaling mutants |  |  |  |
| --- | --- | --- | --- |
| P4-P10 | Total | */* | Recovery |
| *R2<sup>F</sup>* | 386 | 85 | 22% |
| *R2<sup>C</sup>* | 505 | 131 | 26% |
| *R2<sup>PG</sup>* | 475 | 78 | 16% |
| *R2<sup>CPG</sup>* | 396 | 70 | 18% |
| *R2<sup>FCPG</sup>* | 646 | 102 | 16% |
| *R2<sup>KD</sup>* | 38 | 0 | 0 |
| E18.5 | Total | */* | Recovery |
| *R2<sup>F</sup>* | 97 | 22 | 23% |
| *R2<sup>C</sup>* | 53 | 14 | 26% |
| *R2<sup>PG</sup>* | 64 | 14 | 22% |
| *R2<sup>CPG</sup>* | 42 | 10 | 24% |
| *R2<sup>FCPG</sup>* | 87 | 22 | 25% |
| *R2<sup>KD</sup>* | 45 | 0 | - |
| Recovery of *Fgfr2* hemizygous signaling mutants |  |  |  |
| P4-P10 | Total | */- | Recovery |
| *R2<sup>F</sup>* | 37 | 0 | - |
| *R2<sup>C</sup>* | 52 | 10 | 19% |
| *R2<sup>PG</sup>* | 47 | 13 | 27% |
| *R2<sup>CPG</sup>* | 53 | 12 | 22% |
| *R2<sup>FCPG</sup>* | 28 | 1 | 3% |
| P0-P1 | Total | Observed | Expected |
| *R2<sup>F</sup>* | 33 | 4 | 12% |
| *R2<sup>FCPG</sup>* | 25 | 2 | 8% |
| E18.5 | Total | Observed | Expected |
| *R2<sup>F</sup>* | 33 | 5 | 15% |
| *R2<sup>FCPG</sup>* | 31 | 10 | 32% |

**Supplementary Information Table 2:** Craniofacial structures differentially affected in conditional signaling mutants. Micro-CT examination of *Fgfr1*^*F/cKO*^; *Fgfr2*^*cKO/cKO*^ and *Fgfr1*^*FCPG/cKO*^; *Fgfr2*^*cKO/cKO*^ mutant mice skull were to *Fgfr1*^*cKO/cKO*^; *Fgfr2*^*cKO/cKO*^ mutants at E18.5. *Fgfr1*^*FCPG/cKO*^; *Fgfr2*^*cKO/cKO*^ mutant skull show a more severe defect than *Fgfr1*^*F/cKO*^; *Fgfr2*^*cKO/cKO*^ mutants. Several structures are differentially affected in *Fgfr1*^*F/cKO*^; *Fgfr2*^*cKO/cKO*^ and *Fgfr1*^*FCPG/cKO*^; *Fgfr2*^*cKO/cKO*^ mutants. Most severe defects are observed in *Fgfr1*^*cKO/cKO*^; *Fgfr2*^*cKO/cKO*^ mutants.

| *C/CPG<F<FCPG* | *C/CPG<F=FCPG* |
| --- | --- |
| Nasal Cartilage |  |
| Mandible |  |
| Squamosal |  |
| Maxilla |  |
|  | Premaxilla |
|  | Frontal bone |
|  | Zygomatic bone |
|  | Tympanic bulla |
|  | Basisphenoid |
|  | Internal pterygoid process |

**Supplementary Information Table 3:**
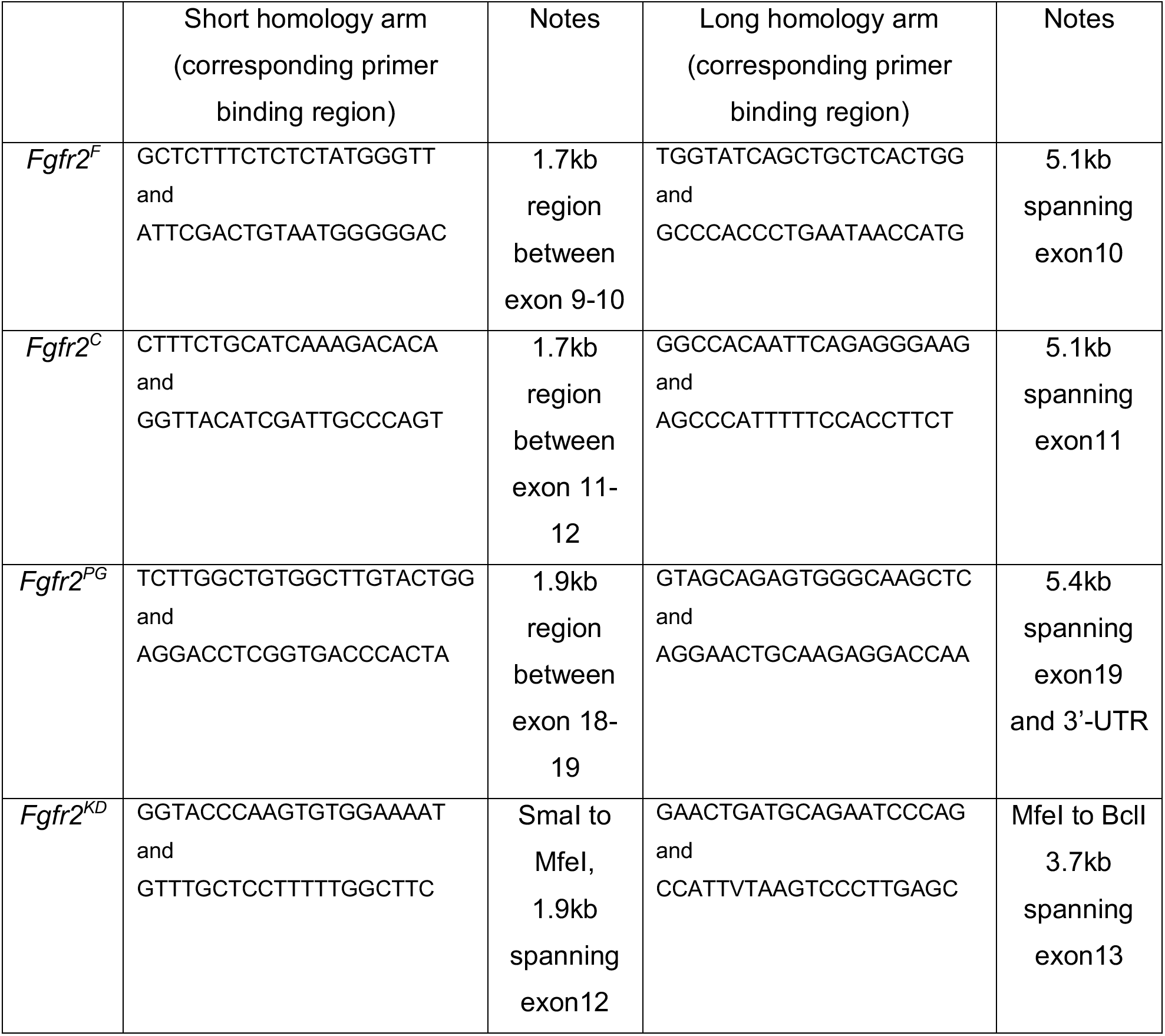
Homology arms used in targeting vectors.

**Supplementary Information Table 4:** Nucleotide and amino-acid substitutions in signaling mutations for *Fgfr2*.

**Supplementary Information Table S4: Nucleotide and amino-acid substitutions in signaling** **mutations for *Fgfr2*.**
| **Mutation** | **Nucleotide substitution** | **Amino acid substitution** | **Notes** |
| --- | --- | --- | --- |
| ***F*** | CTG to GCC<br>AGA to GCA | L424A<br>R426A | Introduces an XmaI site |
| ***C*** | TAT to TTT<br>TTG to CTC | Y466F<br>L468L | Introduces a SacI site |
| ***PG*** | TAC to TTC<br>TAT to TTT | Y769F<br>Y779F | Introduces an EcoRI site |
| ***KD*** | GAAG to AGCT | K517A | Introduces an AluI site |

**Supplementary Information Table 5:**
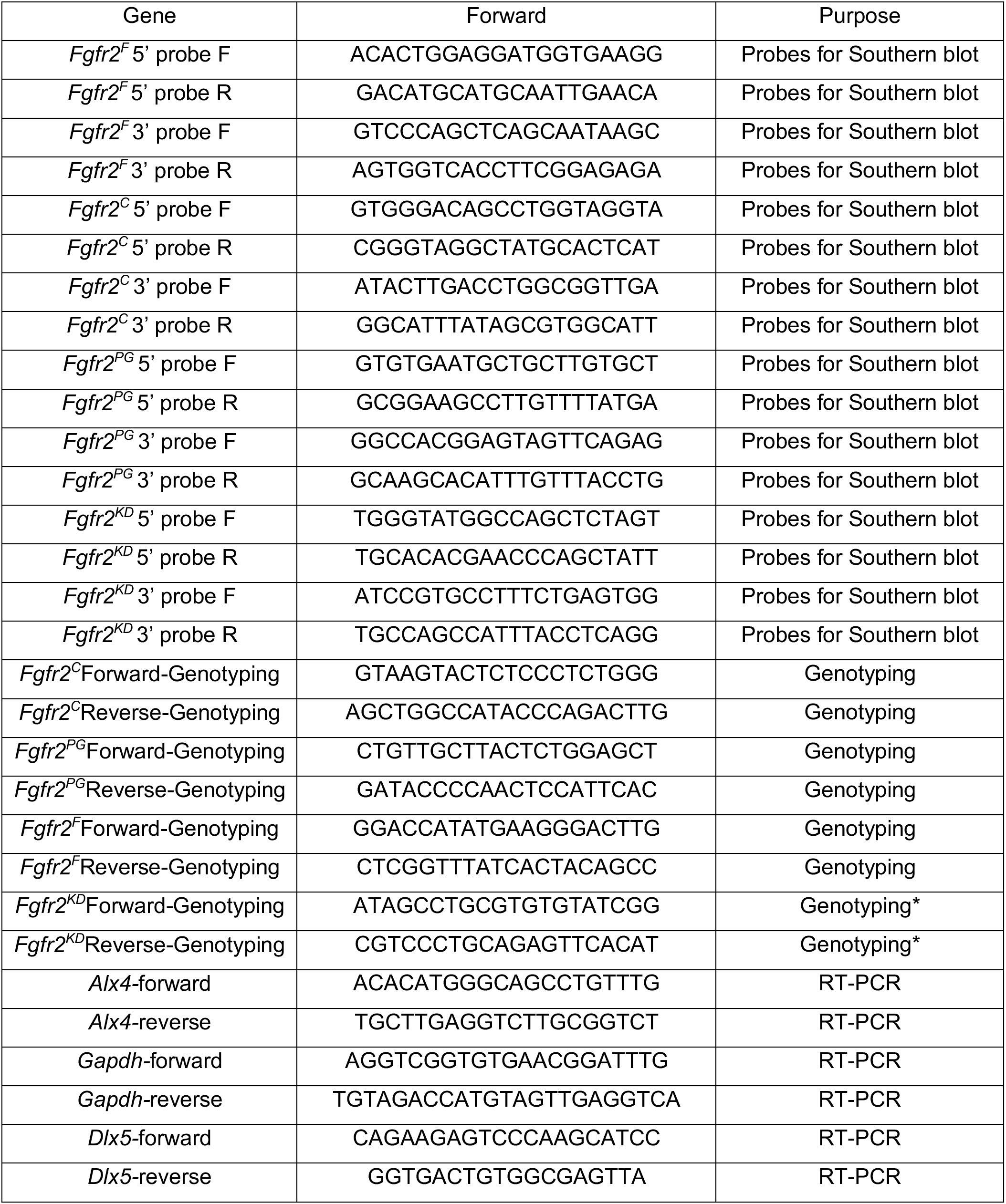
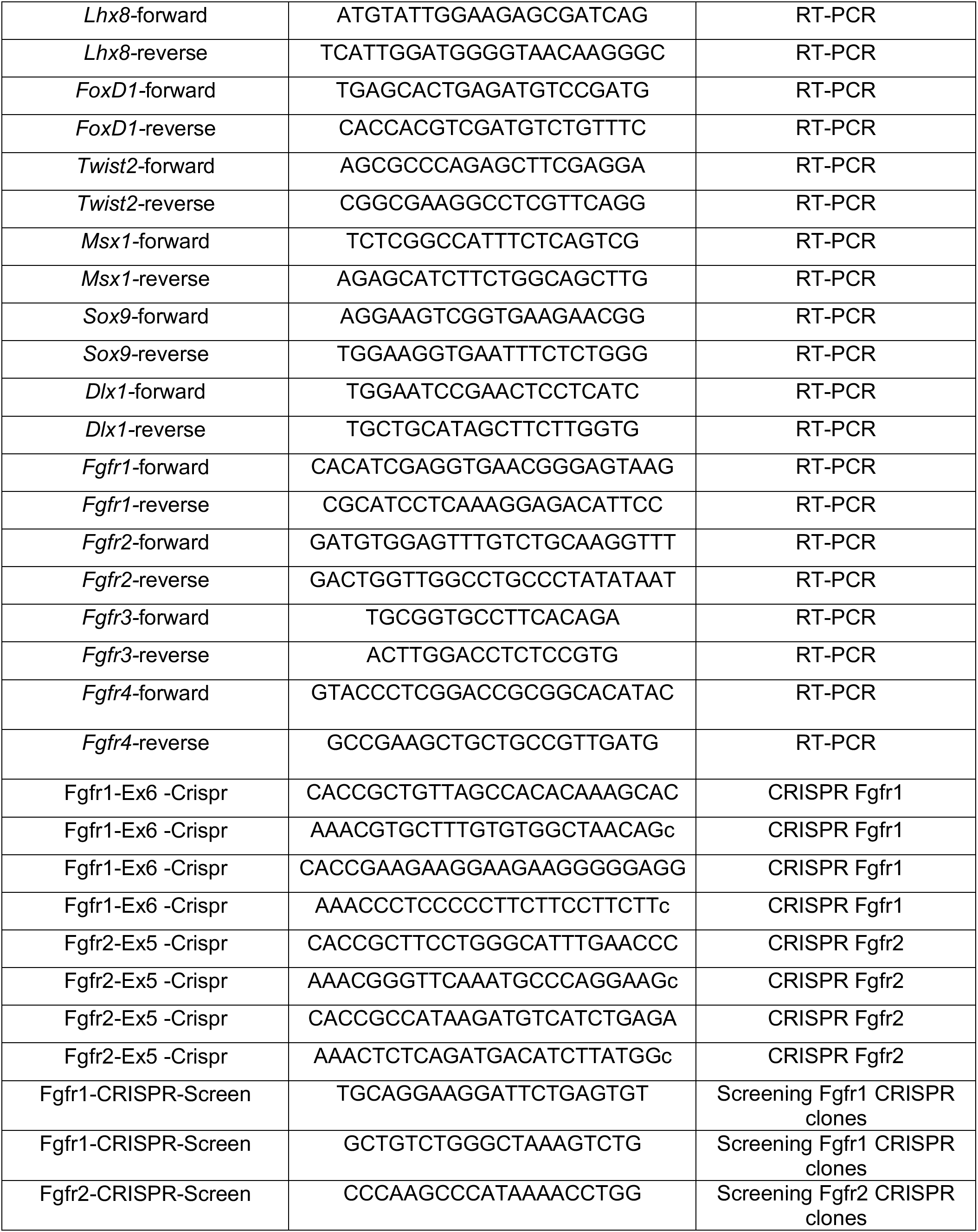
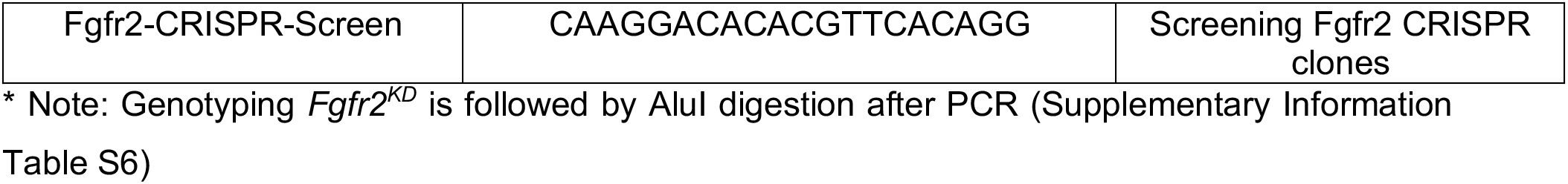
List of oligonucleotides used in this study.

**Supplementary Information Table 6:** Genotyping reactions to identify *Fgfr2* signaling mutations.

**Supplementary Information Table S6: Genotyping reactions to identify *Fgfr2* signaling mutations.**
|  | WT | Mutant |
| --- | --- | --- |
| *Fgfr2<sup>C</sup>* | 553 bp | 655 bp |
| *Fgfr2<sup>PG</sup>* | 377 bp | 611 bp |
| *Fgfr2<sup>F</sup>* | 379 bp | 614 bp |
| *Fgfr2<sup>KD</sup>* | 350 bp | 300 bp |
|  | After Alul digest | After Alul digest |

**Supplementary Information Table 7.**

**Supplementary information Table S7**
|  | Source | Identifier |
| --- | --- | --- |
| **Antibodies** |  |  |
| FGFR2 | Abcam | #ab109372 |
| CRKL | Santa Cruz Biotechnology | #sc-319 |
| FRS2 | Santa Cruz Biotechnology | #sc-8318 |
| FLAG2-M2 | Sigma | #F1804 |
| phospho-p44/42 MAPK<br>Thr202/Tyr204 | Cell Signaling Technology | #9101 |
| p44/42 MAPK | Cell Signaling Technology | #9102 |
| GAPDH | ProteinTech | #60004-1-Ig |
| phospho-AKT Ser473 | Cell Signaling Technology | #4060 |
| AKT | Cell Signaling Technology | #9272 |
| phospho-p38 | Cell signaling Technology | #4511 |
| p38 | Cell signaling Technology | #9212 |
| phospho-PLCγ1 (Y783) | Cell Signaling Technology | #2821 |
| PLCγ1 | Cell Signaling Technology | #2822 |
| phospho-JNK | Cell Signaling Technology | #4671 |
| STAT3α | Cell signaling Technology | #8768 |
| phospho-STAT3 (Y705) | Cell Signaling Technology | #9145 |
| phospho-FAK | Cell Signaling Technology | #3283 |
| FAK | Cell Signaling Technology | #3285 |
| Paxillin | Abcam | #ab32084 |
| IRS2 | Cell Signaling Technology | #3089 |
| β-catenin | Cell Signaling Technology | #8480 |
| Neurofilament antibody | DSHB | #Clone 2H3 |
| EOMES | Abcam | #ab23345 |
| CDX2 | Biogenex | #MU392A-UC |
| GFP | AvesLab | #GFP-1020 |
| SMA | Cell Signaling Technology | #19245 |
| Peroxidase AffiniPure<br>Goat Anti-Mouse IgG<br>(H+L) | Jackson ImmunoResearch<br>Laboratories | #115-035-003 |
| Peroxidase AffiniPure<br>Goat Anti-Rabbit IgG<br>(H+L) | Jackson ImmunoResearch<br>Laboratories | #111-035-003 |
| Donkey anti-Rabbit IgG<br>(H+L), Alexa Fluor 488 | ThermoFisher Scientific | #A-21206 |
| Donkey anti-Rabbit IgG<br>(H+L), Alexa Fluor 546 | ThermoFisher Scientific | #A-10040 |
| Donkey anti-Rabbit IgG<br>(H+L), Alexa Fluor 647 | ThermoFisher Scientific | #A-31573 |
| Donkey anti-Mouse IgG<br>(H+L), Alexa Fluor 488 | ThermoFisher Scientific | #A-21202 |
| Donkey anti- Mouse IgG<br>(H+L), Alexa Fluor 546 | ThermoFisher Scientific | #A-10036 |
| Donkey anti- Mouse IgG<br>(H+L), Alexa Fluor 647 | ThermoFisher Scientific | #A-31571 |
| Alexa Fluor® 488<br>AffiniPure Donkey Anti-Chicken IgY (IgG) (H+L) | Jackson ImmunoResearch<br>Laboratories | #703-545-155 |

## METHODS

### Generation of knock-in mice

Four distinct targeting vectors carrying the *Fgfr2*^*F*^, *Fgfr2*^*C*^, *Fgfr2*^*PG*^, and *Fgfr2*^*KD*^ mutations were generated. The *Fgfr2*^*F*^ targeting vector was generated by cloning a short homology arm (1.7kb region between exon 9-10) and a long homology arm (5.1kb, spanning exon10) into PGKneolox2DTA.2 ^48^. To allow recombineering into SW105 bacteria, the neo cassette was subsequently replaced by PGKEm7neo flanked by FRT sites, which contains both a eukaryotic and a prokaryotic promoter. Similarly, for the *Fgfr2*^*C*^ targeting vector, we cloned a long homology arm (5.3kb region spanning exon11) and a short homology arm (1.7kb region between exon 11-12) into PGKneolox2DTA.2 and used a PGKEm7neo flanked by both FRT and LoxP sites for recombineering. For the *Fgfr2*^*PG*^ targeting vector, we cloned a short homology arm (1.9kb region 5’ of exon19) and a long homology arm (5.4kb spanning exon19 and 3’-UTR) into PGKneolox2DTA.2 and used a PGKEm7neo flanked by FRT sites for recombineering. For *Fgfr2*^*KD*^ targeting vector, we cloned short homology arm (1.9kb SmaI to MfeI, spanning exon 12) and a 3.7kb long homology arm (MfeI to BclI, spanning exon 13) into PGKneolox2DTA.2 ^48^. Details of regions corresponding to homology arms are provided in Supplementary Information Table S3.

For all four alleles, site-directed mutagenesis (SDM) was performed using Phusion polymerase (NEB#M0530). Nucleotide substitutions introduced by SDM in exon10 for *Fgfr2*^*F*^ allele (introduces an XmaI site), exon11 for *Fgfr2*^*C*^ allele (introduces a SacI site), exon 19 for *Fgfr2*^*PG*^ allele (introduces an EcoRI site) and in exon12 for *Fgfr2*^*KD*^ allele (introduces an AluI site) are provided below. All introduced mutations were verified by sequencing. Details of nucleotide substitutions in *Fgfr2* signaling mutations are provided in Supplementary Information Table S4.

The targeting vectors for *Fgfr2*^*F*^ (linearized with NotI), *Fgfr2*^*C*^ (linearized with XhoI), *Fgfr2*^*PG*^ (linearized with NotI) and *Fgfr2*^*KD*^ (linearized with NotI) were electroporated into 129S4 AK7 ES cells. For generating the allelic series of signaling mutations, ES cells were targeted first with the C targeting vector generating *Fgfr2*^+/*C*^ mutant cells. After verifying for correct targeting events, the neo cassette was removed by transient transfection with PGKCrebpA, leaving a single LoxP site behind (Figure 3C). *Fgfr2*^+/*C*^ ES cells were then targeted using the PG targeting vector generating either *Fgfr2*^+/*PG*^ or *Fgfr2*^+/*CPG*^ mutant cells, as determined by breeding of the chimeras to *ROSA26*^*Flpo*^ mice ^49^. After verifying for correct targeting events, the neo cassette was removed by transient transfection with PGKFlpobpA ^50^, leaving both an FRT site and a LoxP site behind (Figure 3C). *Fgfr2*^+/*CPG*^ neo^−^ ES cells were finally targeted with the F targeting vector, resulting in *Fgfr2*^+/*F*^ or *Fgfr2*^+/*FCPG*^ mutant ES cells, as determined by breeding. After verifying for correct targeting events, the neo cassette was removed by transient transfection with Flpe, which is less efficient than Flpo in ES cells ^50^, in order to not recombine sequences between exons 10-18 due to the retention of the FRT site during the generation of the *Fgfr2*^*PG*^ allele. We screened targeting events initially by PCR coupled with restriction digestion to identify incorporation of nucleotide substitutions. Proper targeting was confirmed by Southern blotting using 5′ external and 3′ external probes amplified using the following primer pairs and then an internal probe against Neo. Primers used to generate probes for confirming targeted clones using Southern blots are described in Supplementary Information Table S5.

ES cell chimeras were bred to *Meox2-Cre* or *ROSA26Flpo* deleter mice ^49,51^ maintained on a 129S4 genetic background to remove the neomycin selection cassette and the deleter alleles were subsequently crossed out. Two independent mouse lines were generated from independent ES cell clones for each allele, and phenotypes were confirmed in both lines. The *Fgfr2*^*C*^, *Fgfr2*^*PG*^, *Fgfr2*^*F*^, *Fgfr2*^*FCPG*^ and *Fgfr2*^*KD*^ alleles were maintained on the 129S4 genetic background. *Fgfr2*^*C*^, *Fgfr2*^*PG*^, *Fgfr2*^*F*^ and *Fgfr*2^*KD*^ mice were genotyped using oligonucleotides listed in Supplementary Information Table S5 and S6, with the F, C or PG primers all being able to genotype *Fgfr2*^*FCPG*^ mice.

### Mouse strains

All animal experimentation was conducted according to protocols approved by the Institutional Animal Care and Use Committee of the Icahn School of Medicine at Mount Sinai. *Fgfr1*^*cKO/cKO*^, *Fgfr2*^*cKO/cKO*^, *Fgfr1-GFP* and *Fgfr2-mCherry* were previously described ^18,48^. *Fgfr1* signaling mutations ^7^ are referred to as *Fgfr1*^*C*^, *Fgfr1*^*F*^, *Fgfr1*^*CPG*^ and *Fgfr1*^*FCPG*^. *Tg(Wnt1-cre)11Rth*, *Tg(Wnt1-cre)2Sor*, *Cdkn2a*^+/*tm1Rdp*^, *Gt(ROSA)26Sor*^*tm4(ACTB-tdTomato,-EGFP)Luo*^, and *Bcl2l11*^*tm1.1Ast*^ are referred to in the text as *Wnt1-Cre*, *Wnt1-Cre2*, *Ink, ROSA26^mT/mG^*, and *Bim* respectively ^8,9,52–54^. All lines were maintained on a 129S4 co-isogenic background, except for *Bim* which was crossed into the *Fgfr1/2* deficient backgrounds after only six generations of backcrossing to 129S4.

### *Generation of Fgfr2*^−*Flag3x*^ expression vector and stable 3T3 expression lines

An *Fgfr2* isoform “c” cDNA isoform was PCR amplified from primary MEFs derived from *Fgfr2*^+/+^, *Fgfr2*^*PG/PG*^, *Fgfr2*^*CPG/CPG*^, *Fgfr2*^*F/F*^ and *Fgfr2*^*FCPGFCPG*^ and subsequently digested with HindIII and XhoI. The fragments were cloned in the pcDNA expression vector and sequence verified. Linearized pcDNA-FGFR2 plasmids were transfected in 3T3 cells cultured in DMEM supplemented with 10% calf serum (HyClone Laboratories #SH30072.03) with 50U/mL each penicillin and streptomycin (Gibco #P0781). Stable clones were selected in 500μg/mL G418 (Fischer Scientific #BP6735). 10 clones from each construct (FGFR2^wt^-, FGFR2^PG^-, FGFR2^CPG^-, FGFR2^F^-, or FGFR2^FCPG^-Flag3x) were expanded and assessed for FLAG expression by western blot. Clones expressing high FGFR2-FLAG levels were selected for further analysis.

### Coimmunoprecipitation and Western blotting

Stable 3T3 cells expressing FGFR2^WT^, FGFR2^PG^, FGFR2^CPG^, FGFR2^F^, or FGFR2^FCPG^-Flag3x were serum-starved (0.1% calf serum supplemented DMEM) overnight, stimulated for 15 mins with 50 ng/mL FGF1 (PeproTech, 450-33A) or FGF8b (PeproTech 100-25B) and 5 μg/mL heparin (Sigma #H3149), and lysed in ice-cold NP-40 lysis buffer (20 mM Tris HCL at pH 8, 137 mM NaCl, 10% glycerol, 1% Nonidet [NP-40], 2 mM EDTA, 25 mM β glycerol phosphate, 1 mM Na3VO_4_, 10 mM NaF, 1× cOmplete, EDTA-free Protease Inhibitor Cocktail. 800 μg cell lysates were subsequently used for immunoprecipitation with Anti-FLAG M2 magnetic beads (Sigma #M8823) using the manufacturer’s protocol. We incubated lysates with anti-FLAG M2 magnetic beads overnight at 4°C followed by five washes with lysis buffer, and precipitated proteins were eluted in Laemmli buffer (10% glycerol, 2% SDS, 0.002% bromophenol blue, 0.062M Tris-HCl, pH 6.8) containing 10% β-mercaptoethanol, heated for 5 min at 95°C, separated by SDS-PAGE and analyzed by western blots.

Western blot analysis was performed according to standard protocols using horseradish peroxidase-conjugated secondary antibodies (1:10,000 dilution) developed by chemiluminescent HRP substrate. Primary antibodies were used at the following dilutions for Western blotting: FGFR2 (1:500 dilution), CRKL (1:500 dilution), FRS2 (1:500 dilution), Flag2 M2 (1:500 dilution), phospho-p44/42 MAPK (1:1,000 dilution), p44/42 MAPK (1:1,000 dilution), GAPDH (1:1000 dilution), phospho-AKT (1:1,000 dilution), AKT (1:1,000 dilution), phospho-p38 (1:500 dilution), p38 (1:500 dilution), phospho-PLCγ1 (Y783) (1:200 dilution), PLCγ1 (1:1000 dilution), pJNK (1:500 dilution), phospho-STAT3 (Y705) (1:500 dilution), STAT3α (1:1000 dilution), phospho-FAK (1:1000 dilution), FAK (1:1000 dilution), β-catenin (1:1000 dilution), Paxillin (1:1000 dilution), and IRS2 (Cell Signaling Technology, 3089). For signaling pathways analyzed, the blots were quantified for three independent biological replicates using ImageLab6.0 analysis tool. Ratio of average of mean intensity value of pERK1/2, pAKT, pPLCγ1, pJNK, pSTAT3α, pP38 to GAPDH ± standard deviation was plotted for each pathway.

### Cell derivation and culture conditions

Primary iFNPs were generated by dissecting the maxillary and nasal prominences of E11.5 *Fgfr2*^+/+^; *Ink*^−/−^, *Fgfr2*^*F/F*^; *Ink*^−/−^, *Fgfr2*^*CPG/CPG*^; *Ink*^−/−^, and E9.5 or E11.5 *Fgfr2*^*FCPG/FCPG*^; *Ink*^−/−^ embryos in PBS. The tissue was disassociated with 0.125% Trypsin-EDTA and cultured in DMEM supplemented with 20% FBS, 50 U/mL each penicillin and streptomycin on fibronectin coated plates (0.5μg/cm^2^). Cells were subsequently split 1:5 through for at least 5 passages before immortalized cell lines were obtained. Cells were allowed to grow until sub-confluent. All experiments were performed between passage 15 and 25. We used PX459 V2.0 vector (Addgene plasmid # 62988) to CRISPR out either *Fgfr1* or *Fgfr2* and create *Fgfr1* null, *Fgfr2* null, or *Fgfr1*: *Fgfr2* double null cells. gRNA sequences for *Fgfr1* and *Fgfr2* were selected using CHOPCHOP gRNA design web tool and were cloned using the oligonucleotides (Supplementary Information Table S5), as previously described ^55^. Plasmids were transfected in respective iFNP cells cultured in DMEM (Sigma #5796) supplemented with 10% calf serum with 50 U/mL each penicillin and streptomycin. Stable clones were selected in 5μg/mL Puromycin (Sigma #P8833). Clones were verified (homozygous deletion of exon6 for *Fgfr1* and deletion exon5 for *Fgfr2* which also introduces a frameshift mutation) using PCR (Supplementary Information Table S5). Primary MEFs were derived from E12.5 wild type mice embryos. Embryos were eviscerated and after removing the head, remaining tissue was chopped into 1 cm pieces and incubated in (0.25%) 1 mL trypsin-EDTA (ThermoFisher # 25200-072) for 30 mins with intermittent shaking. 10 mL DMEM 10% calf serum was added and the mixture was allowed to pass through a cell-strainer. Cells collected from each embryo was plated in 0.2% gelatin (Sigma #G1393) coated 15cm plate.

### Skeletal preparations

Embryos at E14.5, E16.5 or E18.5 embryos were skinned, eviscerated, fixed in 95% ethanol overnight, and stained (0.015% Alcian blue, 0.005% Alizarin red, 5% glacial acetic acid, in 70% ethanol) overnight at 37°C. Skeletons were then cleared in 1% KOH and transferred to decreasing concentrations of KOH in increasing concentrations of glycerol until clear.

### Acetocarmine and hematoxylin and eosin staining

Freshly harvested tissue was fixed in 4% PFA at 4°C overnight followed by dehydration in 70% ethanol. For acetocarmine staining, tissues were incubated in 0.5% aceto-carmine (0.5 g carmine stain (Sigma #C1022) dissolved in 100 ml boiling 45% acetic acid for 15 minutes), followed by de-staining in 70% ethanol for 1 minute and 1% acid alcohol (1% HCl in 70% ethanol) for 2 minutes and 5% acid alcohol (5% HCl in 70% ethanol) for 1 minute. For hematoxylin and eosin staining, freshly harvested tissues were dissected in PBS, and fixed in 4% PFA followed by dehydration through a graded ethanol series, and embedded in paraffin. 5μm sections were cut. After deparaffinization and rehydration, sections were stained with Harris modified hematoxylin (Sigma #HHS16) followed by a 10 second wash in acid-alcohol (1% v/v HCl in 70% EtOH), followed by counterstaining with 1% eosinY (Sigma #17372-87-1). Tissues were washed and mounted with Permount (ThermoFisher Scientific #SP15100).

### Scratch assays

Cells were seeded onto glass coverslips coated with 5 μg/mL human plasma fibronectin purified protein (Millipore Sigma #FC010). At ~90–100% confluency, cells were scratched with a P1000 pipet tip, washed with PBS and incubated in fresh medium containing either 0.1% FBS, 10% FBS, 50 ng/mL FGF1 and 5 μg/mL heparin or 10 ng/mL PDGF-AA (R&D Systems #1055-AA-050) supplemented DMEM for 12 hrs.

### Immunofluorescence and Antibodies

For immunostaining whole mount embryos were fixed in 4% paraformaldehyde solution (PFA) in PBS overnight and washed with PBS five times, permeabilized with 0.5% Triton X100 in PBS for 30 min and blocked in 2% blocking serum for 2h at room temperature. Primary anti-neurofilament antibody (Clone 2H3, DSHB) was used at a 1:20 dilution in 1% donkey serum in PBST; embryos were incubated overnight at 4°C. The next day, embryos were washed 4 times in PBST and incubated with anti-mouse HRP-conjugated secondary antibodies at a 1:1000 dilution for 4hrs at room temperature followed by washing in PBST 4 times and signal was developed using ImmPACTDAB kit (VectorLabs #SK-4105). For whole-mount immunofluorescence at E7.5, embryos were fixed overnight in 4:1 methanol:DMSO. Primary antibodies for Eomes (1:100 dilution) and Cdx2 (1:100 dilution) were used. For immunostaining cells, cells were fixed for 10 mins in 4% PFA in PBS at room temperature. Cells/ tissues were subsequently processed for immunofluorescence analysis as detailed above using anti-paxillin primary antibody (1:250 dilution) with Alexa647 conjugated phalloidin (1:40 dilution). For immunofluorescence on sections, antibodies for GFP (1:100 dilution), mCherry (1:100 dilution) and SMA (1:100 dilution) was used. Supplementary Information Table S7 provides more information on antibodies used in this study. Embryos were stained with DAPI following fixation as previously described ^56^. Cells / tissues were photographed using an Olympus DP71 digital camera fitted onto an Olympus BX51 fluorescence microscope, Leica SP5 confocal or a Hamamatsu C11440 camera fitted to a Zeiss Observer Z1 microscope. Epifluorescence was imaged in Zeiss Axioplan fitted to ProgRes CT3 camera.

### In situ hybridization

Labeled antisense-RNA probes were synthesized for *Alx3, Msx1, Six3, Nkx2.1, Fgf8, Shh, Col2a1, Col10a1* and *Meox1*. Digoxigenin-labeled anti-sense probes were generated as described, and mRNA in situ hybridization on paraffin sections for chromogenic detection was performed using standard protocols.

### Micro-CT imaging

Micro-CT imaging of the skulls were performed using a SkyScan 1172 scanner (Bruker, Kontich, Belgium). The mouse heads were dissected and fixed in 10% neutral buffered formalin and washed and stored in PBS at 4 °C. The skull bones were scanned with settings of 50 kV, 500 μA, 10 μm pixel resolution, 0.3° rotation steps, and 4 frames average imaging with a 0.5-mm Al filter at Micro-CT Core, School of Dentistry, NYU, New York. The acquired X-ray projections were reconstructed using the Imaris software (Oxford Instruments).

### Cell proliferation assay

For EdU labeling in mice, pregnant females were injected intraperitoneally with 100 mg/kg body weight of EdU. EdU detection was carried out as per manufacturer’s instruction for Click-iT EdU Cell Proliferation Kit.

### TUNEL assay

Sections were deparaffinized and were rehydrated in PBS, followed by post-fixation in 4% PFA. In Situ Cell Death Detection Kit, TMR red user protocol was used to detect cell death.

### RT-qPCR

Cells were lysed, and mRNA was extracted according to Qiagen RNeasy kit standard protocol. cDNA was synthesized using a 2:1 ratio of random primers to Oligo(dT) with SuperScript IV RT (Invitrogen). qPCR was performed with PerfeCTa SYBR Green FastMix for iQ (Quanta Biosciences) with Bio-Rad iQ5 multicolor real-time PCR detection system and analyzed with Bio-Rad iQ5 optical system software (version 2.0). Cycling conditions were as follows: step 1, 3 min at 95°C; step 2, 10 sec at 95°C; step 3, 30 sec at 60°C; and repeat steps 2 and 3 for 40 cycles. Proper amplification was confirmed using a melting curve and by running samples on a gel to ensure that the correct size band was obtained. Graphs were made using Microsoft Excel and Prism. Primer sequence for respective genes used for RT-qPCR analysis is listed below.

### β-Catenin Quantification

Cells were stained and imaged using a Leica SP5 confocal microscope under identical conditions. Stacks were then background subtracted using a 100px rolling ball function in ImageJ. The average pixel intensity along cell-cell junctions was measured in a single z-plane per junction using a 15-pixel line width in ImageJ for the EGFP and β-catenin channels.

### Quantification and statistical analysis

Statistical analysis was performed using GraphPad Prism6.0 and Microsoft Excel. Values are presented as mean ± standard deviation. The statistical significance was determined using a student *t*-test with Holm-Sidak method.

## References

1 Lemmon, M. A. & Schlessinger, J. Cell signaling by receptor tyrosine kinases. Cell 141, 1117–1134, doi:10.1016/j.cell.2010.06.011 (2010).

2 Zinkle, A. & Mohammadi, M. A threshold model for receptor tyrosine kinase signaling specificity and cell fate determination. F1000Res 7, doi:10.12688/f1000research.14143.1 (2018).

3 Li, P. & Elowitz, M. B. Communication codes in developmental signaling pathways. Development 146, doi:10.1242/dev.170977 (2019).

4 Vasudevan, H. N., Mazot, P., He, F. & Soriano, P. Receptor tyrosine kinases modulate distinct transcriptional programs by differential usage of intracellular pathways. Elife 4, doi:10.7554/eLife.07186 (2015).

5 Brewer, J. R., Mazot, P. & Soriano, P. Genetic insights into the mechanisms of Fgf signaling. Genes Dev 30, 751–771, doi:10.1101/gad.277137.115 (2016).

6 Lanner, F. & Rossant, J. The role of FGF/Erk signaling in pluripotent cells. Development 137, 3351–3360 (2010).

7 Brewer, J. R., Molotkov, A., Mazot, P., Hoch, R. V. & Soriano, P. Fgfr1 regulates development through the combinatorial use of signaling proteins. Genes Dev 29, 1863–1874, doi:10.1101/gad.264994.115 (2015).

8 Danielian, P. S., Muccino, D., Rowitch, D. H., Michael, S. K. & McMahon, A. P. Modification of gene activity in mouse embryos in utero by a tamoxifen-inducible form of Cre recombinase. Curr Biol 8, 1323–1326, doi:10.1016/s0960-9822(07)00562-3 (1998).

9 Lewis, A. E., Vasudevan, H. N., O’Neill, A. K., Soriano, P. & Bush, J. O. The widely used Wnt1-Cre transgene causes developmental phenotypes by ectopic activation of Wnt signaling. Dev Biol 379, 229–234, doi:10.1016/j.ydbio.2013.04.026 (2013).

10 Partanen, J., Schwartz, L. & Rossant, J. Opposite phenotypes of hypomorphic and Y766 phosphorylation site mutations reveal a function for Fgfr1 in anteroposterior patterning of mouse embryos. Genes Dev 12, 2332–2344 (1998).

11 De Moerlooze, L. et al. An important role for the IIIb isoform of fibroblast growth factor receptor 2 (FGFR2) in mesenchymal-epithelial signalling during mouse organogenesis. Development 127, 483–492 (2000).

12 Youle, R. J. & Strasser, A. The BCL-2 protein family: opposing activities that mediate cell death. Nat Rev Mol Cell Biol 9, 47–59, doi:10.1038/nrm2308 (2008).

13 Chipuk, J. E. & Green, D. R. How do BCL-2 proteins induce mitochondrial outer membrane permeabilization? Trends Cell Biol 18, 157–164, doi:10.1016/j.tcb.2008.01.007 (2008).

14 Czabotar, P. E., Lessene, G., Strasser, A. & Adams, J. M. Control of apoptosis by the BCL-2 protein family: implications for physiology and therapy. Nat Rev Mol Cell Biol 15, 49–63, doi:10.1038/nrm3722 (2014).

15 Grabow, S. et al. Subtle Changes in the Levels of BCL-2 Proteins Cause Severe Craniofacial Abnormalities. Cell Rep 24, 3285–3295 e3284, doi:10.1016/j.celrep.2018.08.048 (2018).

16 Clybouw, C. et al. Alternative splicing of Bim and Erk-mediated Bim(EL) phosphorylation are dispensable for hematopoietic homeostasis in vivo. Cell Death Differ 19, 1060–1068, doi:10.1038/cdd.2011.198 (2012).

17 Lei, K. & Davis, R. J. JNK phosphorylation of Bim-related members of the Bcl2 family induces Bax-dependent apoptosis. Proc Natl Acad Sci U S A 100, 2432–2437, doi:10.1073/pnas.0438011100 (2003).

18 Molotkov, A., Mazot, P., Brewer, J. R., Cinalli, R. M. & Soriano, P. Distinct Requirements for Fgfr1 and Fgfr2 in Primitive Endoderm Development and Exit from Pluripotency. Developmental Cell 41, 511–526 (2017).

19 Yu, K. et al. Conditional inactivation of FGF receptor 2 reveals an essential role for FGF signaling in the regulation of osteoblast function and bone growth. Development 130, 3063–3074 (2003).

20 Xu, X. et al. Fibroblast growth factor receptor 2 (FGFR2)-mediated reciprocal regulation loop between FGF8 and FGF10 is essential for limb induction. Development 125, 753–765 (1998).

21 Garg, A. et al. Alx4 relays sequential FGF signaling to induce lacrimal gland morphogenesis. PLoS Genet 13, e1007047, doi:10.1371/journal.pgen.1007047 (2017).

22 Steinberg, Z. et al. FGFR2b signaling regulates ex vivo submandibular gland epithelial cell proliferation and branching morphogenesis. Development 132, 1223–1234, doi:10.1242/dev.01690 (2005).

23 Francavilla, C. et al. Functional proteomics defines the molecular switch underlying FGF receptor trafficking and cellular outputs. Mol Cell 51, 707–722, doi:10.1016/j.molcel.2013.08.002 (2013).

24 Kurowski, A., Molotkov, A. & Soriano, P. FGFR1 regulates trophectoderm development and facilitates blastocyst implantation. Dev Biol 446, 94–101, doi:10.1016/j.ydbio.2018.12.008 (2019).

25 Fantauzzo, K. A. & Soriano, P. Generation of an Immortalized Mouse Embryonic Palatal Mesenchyme Cell Line. PLoS ONE 12, e0179078 (2017).

26 Hanks, S. K., Quinn, A. M. & Hunter, T. The protein kinase family: conserved features and deduced phylogeny of the catalytic domains. Science 241, 42–52, doi:10.1126/science.3291115 (1988).

27 Bellot, F. et al. Ligand-induced transphosphorylation between different FGF receptors. EMBO J 10, 2849–2854 (1991).

28 Ueno, H., Gunn, M., Dell, K., Tseng, A., Jr. & Williams, L. A truncated form of fibroblast growth factor receptor 1 inhibits signal transduction by multiple types of fibroblast growth factor receptor. J Biol Chem 267, 1470–1476 (1992).

29 Meyer, M. et al. FGF receptors 1 and 2 are key regulators of keratinocyte migration in vitro and in wounded skin. J Cell Sci 125, 5690–5701, doi:10.1242/jcs.108167 (2012).

30 Rasouli, S. J. et al. The flow responsive transcription factor Klf2 is required for myocardial wall integrity by modulating Fgf signaling. Elife 7, doi:10.7554/eLife.38889 (2018).

31 Sun, J. & Stathopoulos, A. FGF controls epithelial-mesenchymal transitions during gastrulation by regulating cell division and apicobasal polarity. Development 145, doi:10.1242/dev.161927 (2018).

32 Yayon, A., Klagsbrun, M., Esko, J. D., Leder, P. & Ornitz, D. M. Cell surface, heparin-like molecules are required for binding of basic fibroblast growth factor to its high affinity receptor. Cell 64, 841–848, doi:10.1016/0092-8674(91)90512-w (1991).

33 Rapraeger, A. C., Krufka, A. & Olwin, B. B. Requirement of heparan sulfate for bFGF-mediated fibroblast growth and myoblast differentiation. Science 252, 1705–1708, doi:10.1126/science.1646484 (1991).

34 Endo, Y., Ishiwata-Endo, H. & Yamada, K. M. Extracellular matrix protein anosmin promotes neural crest formation and regulates FGF, BMP, and WNT activities. Dev Cell 23, 305–316, doi:10.1016/j.devcel.2012.07.006 (2012).

35 Geiger, B. & Yamada, K. M. Molecular architecture and function of matrix adhesions. Cold Spring Harb Perspect Biol 3, doi:10.1101/cshperspect.a005033 (2011).

36 McQuade, K. J., Beauvais, D. M., Burbach, B. J. & Rapraeger, A. C. Syndecan-1 regulates alphavbeta5 integrin activity in B82L fibroblasts. J Cell Sci 119, 2445–2456, doi:10.1242/jcs.02970 (2006).

37 Moser, M., Legate, K. R., Zent, R. & Fassler, R. The tail of integrins, talin, and kindlins. Science 324, 895–899, doi:10.1126/science.1163865 (2009).

38 Frisch, S. M. & Francis, H. Disruption of epithelial cell-matrix interactions induces apoptosis. J Cell Biol 124, 619–626, doi:10.1083/jcb.124.4.619 (1994).

39 Mailleux, A. A. et al. BIM regulates apoptosis during mammary ductal morphogenesis, and its absence reveals alternative cell death mechanisms. Dev Cell 12, 221–234, doi:10.1016/j.devcel.2006.12.003 (2007).

40 Borges, E., Jan, Y. & Ruoslahti, E. Platelet-derived growth factor receptor beta and vascular endothelial growth factor receptor 2 bind to the beta 3 integrin through its extracellular domain. J Biol Chem 275, 39867–39873, doi:10.1074/jbc.M007040200 (2000).

41 Klinghoffer, R. A., Hamilton, T. G., Hoch, R. & Soriano, P. An allelic series at the PDGFalphaR locus indicates unequal contributions of distinct signaling pathways during development. Dev Cell 2, 103–113 (2002).

42 Tallquist, M. D., French, W. J. & Soriano, P. Additive effects of PDGF receptor beta signaling pathways in vascular smooth muscle cell development. PLoS Biol 1, E52, doi:10.1371/journal.pbio.0000052 (2003).

43 Kon, E. et al. N-cadherin-regulated FGFR ubiquitination and degradation control mammalian neocortical projection neuron migration. Elife 8, doi:10.7554/eLife.47673 (2019).

44 Scarpa, E. et al. Cadherin Switch during EMT in Neural Crest Cells Leads to Contact Inhibition of Locomotion via Repolarization of Forces. Dev Cell 34, 421–434, doi:10.1016/j.devcel.2015.06.012 (2015).

45 Ciruna, B. & Rossant, J. FGF signaling regulates mesoderm cell fate specification and morphogenetic movement at the primitive streak. Dev Cell 1, 37–49 (2001).

46 Sun, X., Meyers, E. N., Lewandoski, M. & Martin, G. R. Targeted disruption of Fgf8 causes failure of cell migration in the gastrulating mouse embryo. Genes Dev 13, 1834–1846, doi:10.1101/gad.13.14.1834 (1999).

47 Nieto, M. A., Huang, R. Y., Jackson, R. A. & Thiery, J. P. Emt: 2016. Cell 166, 21–45, doi:10.1016/j.cell.2016.06.028 (2016).

48 Hoch, R. V. & Soriano, P. Context-specific requirements for Fgfr1 signaling through Frs2 and Frs3 during mouse development. Development 133, 663–673, doi:10.1242/dev.02242 (2006).

49 Raymond, C. S. & Soriano, P. ROSA26Flpo deleter mice promote efficient inversion of conditional gene traps in vivo. Genesis 48, 603–606, doi:10.1002/dvg.20659 (2010).

50 Raymond, C. S. & Soriano, P. High-efficiency FLP and PhiC31 site-specific recombination in mammalian cells. PLoS One 2, e162, doi:10.1371/journal.pone.0000162 (2007).

51 Tallquist, M. D. & Soriano, P. Epiblast-restricted Cre expression in MORE mice: a tool to distinguish embryonic vs. extra-embryonic gene function. Genesis 26, 113–115 (2000).

52 Bouillet, P. et al. Proapoptotic Bcl-2 relative Bim required for certain apoptotic responses, leukocyte homeostasis, and to preclude autoimmunity. Science 286, 1735–1738, doi:10.1126/science.286.5445.1735 (1999).

53 Muzumdar, M. D., Tasic, B., Miyamichi, K., Li, L. & Luo, L. A global double-fluorescent Cre reporter mouse. Genesis 45, 593–605, doi:10.1002/dvg.20335 (2007).

54 Serrano, M. et al. Role of the INK4a locus in tumor suppression and cell mortality. Cell 85, 27–37, doi:10.1016/s0092-8674(00)81079-x (1996).

55 Ran, F. A. et al. Genome engineering using the CRISPR-Cas9 system. Nat Protoc 8, 2281–2308, doi:10.1038/nprot.2013.143 (2013).

56 Sandell, L. L., Kurosaka, H. & Trainor, P. A. Whole mount nuclear fluorescent imaging: convenient documentation of embryo morphology. Genesis 50, 844–850, doi:10.1002/dvg.22344 (2012).

